# Two Specialized Intrinsic Cardiac Neuron Types Safeguard Heart Homeostasis and Stress Resilience

**DOI:** 10.1101/2025.06.13.659555

**Authors:** Qian J. Xu, Marissa C. Applegate, I-Uen Hsu, Rachel P. Kogan, Omar A. Hafez, Ruiqi L. Wang, Pam E. Rios Coronado, Lawrence H. Young, Xing Zeng, Le Zhang, Rui B. Chang

**Affiliations:** Department of Neuroscience, Yale University School of Medicine, New Haven, CT 06520, USA; Department of Cellular and Molecular Physiology, Yale University School of Medicine, New Haven, CT 06520, USA; Interdepartmental Neuroscience Program, Yale University School of Medicine, New Haven, CT 06520, USA; M.D.-Ph.D. Program, Yale University School of Medicine, New Haven, CT 06520, USA; Section of Cardiovascular Medicine, Department of Internal Medicine, Yale University School of Medicine, New Haven, CT 06520, USA; Department of Physiology, University of Texas Southwestern Medical Centre, Dallas, TX 75235, USA; Department of Neurology, Yale University School of Medicine, New Haven, CT 06520, USA

## Abstract

The intrinsic cardiac nervous system (ICNS), often referred to as “the little brain on the heart”, plays a central role in heart-brain communication and is increasingly recognized as both a contributor to cardiac disorders, including atrial fibrillation, heart failure, and sudden cardiac death, and a growing target for therapeutic intervention. Despite its clinical relevance, the molecular and functional organization of the ICNS remains poorly understood. Using single-cell transcriptomics, high-resolution imaging, and cell-specific genetic tools in mice, we identified two molecularly distinct intrinsic cardiac neuron (ICN) subtypes marked by *Npy* and *Ddah1*, each exhibiting unique anatomical innervation patterns. *Npy⁺* ICNs function as parasympathetic postganglionic neurons essential for maintaining coronary perfusion and cardiac homeostasis during routine physiological activity. In contrast, *Ddah1*⁺ ICNs are crucial for preserving cardiac electrical stability and preventing sudden death under extreme stress. These findings uncover specialized ICNS pathways that support cardiac homeostasis and resilience, providing a foundation for developing targeted neurocardiac therapies for autonomic dysfunction in human heart disease.

## Introduction

The heart, as one of the body’s vital organs, must operate continuously and adaptively to sustain life. This demand requires precise regulation of cardiac homeostasis under both routine and extreme physiological conditions. The brain exerts tight control over heart function through the autonomic nervous system^1–4^, a complex network of afferent and efferent pathways that modulate heart rhythm, cardiac output, and vascular tone. Traditionally, studies of neurocardiac regulation have focused primarily on extrinsic autonomic pathways, particularly vagal and spinal sensory afferents, as well as sympathetic and parasympathetic efferents. While these brain-to-heart circuits are well established as key regulators of cardiovascular physiology and homeostasis^1–10^, the discovery of the intrinsic cardiac nervous system (ICNS), a complex neural network embedded within the heart, has revealed a more intricate framework of neurocardiac control^11–14^.

Anatomically situated at the interface between extrinsic autonomic inputs and diverse cardiac cell types, such as pacemaker cells and cardiomyocytes, the ICNS is uniquely positioned to integrate central and local signals. Once regarded as a passive parasympathetic relay, the ICNS is now thought to function similarly to the enteric nervous system (ENS), comprising local populations of sensory neurons, interneurons, and motor neurons capable of monitoring, integrating, and responding to cardiac activity^14^. Consistent with this model, recent studies have identified a broad range of neurochemical phenotypes within the ICNS^11,15–20^, leading to its characterization as the “little brain on the heart”. Clinically, the ICNS has been implicated in a wide spectrum of cardiovascular disorders, including atrial and ventricular fibrillation, myocardial ischemia, heart failure, and sudden cardiac death^2,11,12,21–24^, underscoring its relevance as a therapeutic target. Indeed, ICNS-targeted interventions have already been applied in the treatment of atrial fibrillation and reflex vasovagal syncope^25–27^. However, classical neuromodulatory approaches, such as electrical, pharmacological, and surgical methods, lack cell-type specificity. This limitation not only impedes mechanistic investigation of ICNS function but also contributes to inconsistent and sometimes contradictory outcomes in clinical settings^27–30^. Despite its central role in cardiac pathophysiology, the molecular identity, structural organization, and functional specialization of intrinsic cardiac neuron (ICN) subtypes remain poorly defined, posing a significant barrier to the development of targeted and effective therapies for heart disease.

To resolve the molecular and functional complexity of the ICNS, we integrated single-cell transcriptomics, high-resolution volumetric imaging, genetic labeling, and cell-specific neuromodulation in mice. This strategy revealed two transcriptionally and anatomically distinct ICN subtypes, defined by expression of *Npy* and *Ddah1*. These subtypes exhibit divergent roles in regulating cardiovascular stability: *Npy⁺* neurons function as parasympathetic postganglionic cells essential for maintaining coronary perfusion and cardiac homeostasis under baseline conditions, whereas *Ddah1⁺* neurons are required to preserve electrical stability and prevent stress-induced cardiac collapse. Together, our findings uncover a division of labor within the ICNS and offer a blueprint for developing cell type-targeted neuromodulatory therapies for heart disease.

## Results

### *Npy* and *Ddah1* differentially label two primary subtypes of ICNs

Understanding the molecular architecture of the ICNS is crucial for mapping its neural circuits and associated physiological functions. ICNs are rare cells on the heart, with only ∼1,000 neurons in the mouse ICNS^31^. To fluorescently label all ICNs, we generated genetically engineered Baf53b-Cre; lox-L10-GFP mice (designated Baf53b^L10-GFP^)^32^ by crossing Baf53b-Cre mice, which exhibit pan-neuronal Cre activity^33^, with mice harboring a Cre-dependent L10-GFP allele (lox-L10-GFP)^34,35^ (Figure 1A). As expected, we observed fluorescently labeled ICNs stereotypically distributed across several anterior and posterior ganglia on the heart atria^32^.

**Figure 1.**
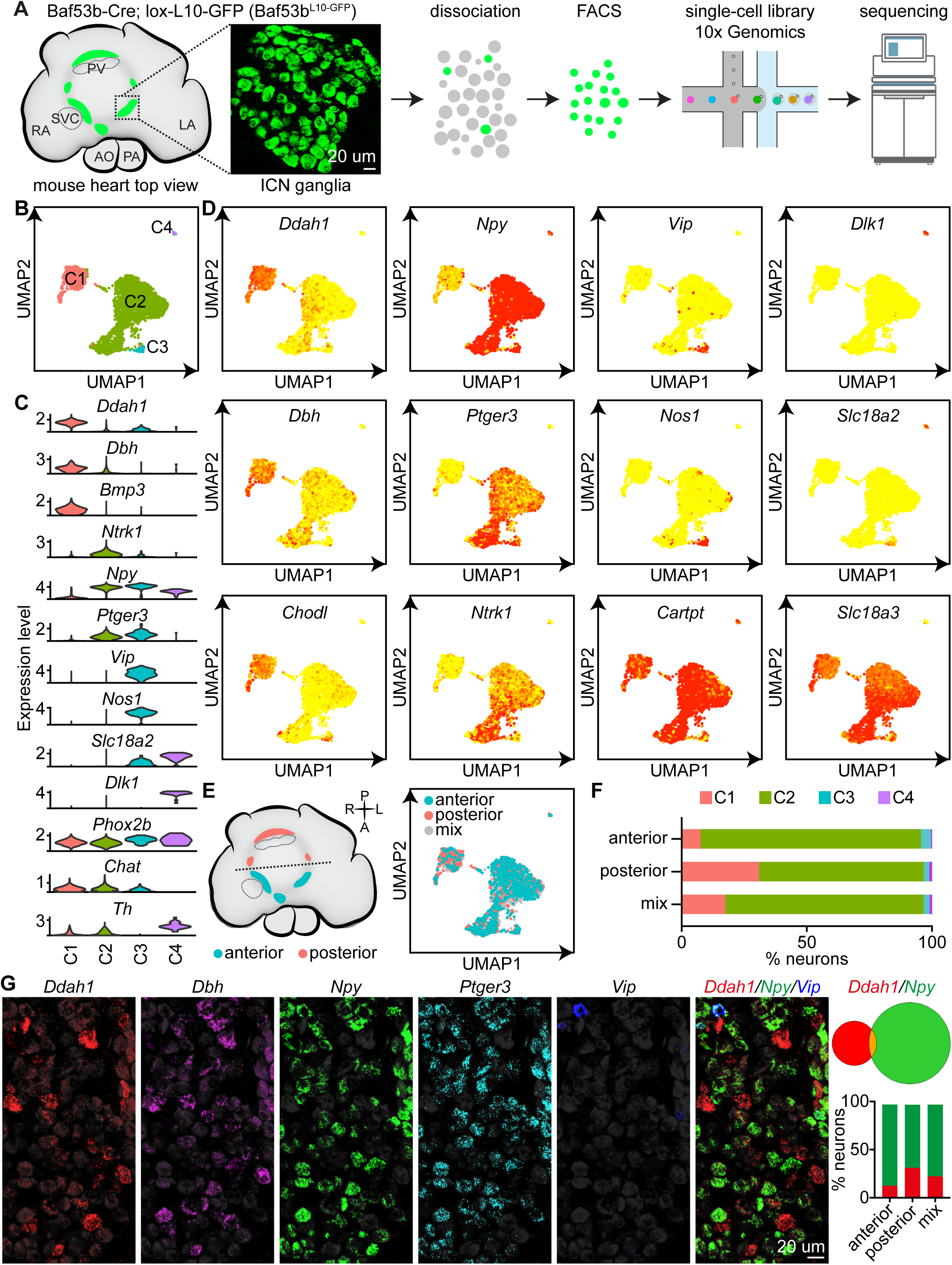
Molecular architecture of the ICNS reveals two major ICN subtypes. (A) Schematic illustration of single cell RNA-sequencing workflow. ICNs (green) labeled in Baf53b^L10-GFP^ mice were acutely isolated from the heart, purified by fluorescence-activated cell sorting (FACS), and subjected to single cell RNA-sequencing using the 10x Genomics platform. RA, right atrium; LA, left atrium; AO, aorta; PA, pulmonary artery; PV, pulmonary vein; SVC, superior vena cava. (B) UMAP plot of 2,870 ICNs showing four major clusters (C1-C4, color coded). (C) Violin plots showing expression of selected marker genes across ICN subtypes. (D) UMAP plots from (B) colored by expression of the indicated marker genes. (E) Schematic illustration (left) and UMAP plot (right) of ICNs sequenced from anterior (cyan), posterior (coral), or mixed (grey) ganglia. (F) Percentage of ICNs in the four subtypes (colors as indicated), grouped by anatomical origin. (G) RNAscope HiPlex Assay for the indicated genes, showing that *Ddah1* and *Npy* are expressed in largely non-overlapping ICNs. ICNs expressing *Ddah1* (red), *Npy* (green) or both (orange) were quantified, and the proportions of *Ddah1*^+^ and *Npy*^+^ ICNs from anterior, posterior, and mixed ganglia were plotted. Scale bars, 20 μm. See also Figures S1 and S2.

ICNs acutely isolated from anterior, posterior, and mixed regions of adult Baf53b^L10-GFP^ mice were purified using Fluorescence-Activated Cell Sorting (FACS) and subjected to single-cell RNA sequencing using the 10x Genomics platform (Figure 1A). A total of 2,870 ICNs were analyzed and unbiasedly grouped into four clusters based on differentially expressed genes (DEGs) using Seurat^36^, and visualized using Uniform Manifold Approximation and Projection (UMAP)^37^ (Figures 1B-D and S1A). As previously reported^34,35,38–40^, all ICNs are *Phox2b^+^* and most of them are cholinergic (*Slc18a3*, 98.2%; *Chat*, 76.6%) and *Cartpt*^+^ (96.9%). While some ICNs are *Th^+^* (65.5%), they do not express classical transporters for norepinephrine (*Slc6a2*, 0.2%), dopamine (*Slc6a3*, 0.9%), or vesicular monoamines (*Slc18a2*, 3.6%), suggesting a lack of adrenergic function. ICNs also lack common sensory markers such as *Piezo1* (0.5%), *Piezo2* (0.1%), *Trpv1* (0.2%), *Tac1* (6.8%), and *Calca* (1.7%), or machineries for glutamate and GABA transmission, indicating that they are less heterogeneous than gut-intrinsic enteric neurons^41^. Among the four clusters, *Ddah1* and *Npy* distinctly label two primary types of ICNs (Figure 1C-D) while Cluster 3 (*Vip*^+^/*Nos1*^+^) and Cluster 4 (*Dlk1*^+^/*Slc18a2*^+^) only represent very small fractions of the ICNS (2.4 and 1.0% respectively). *Dbh* and *Chodl* were selectively expressed in *Ddah1*^+^ ICNs whereas *Npy*^+^ ICNs co-expressed *Ptger3* and *Ntrk1* (Figure 1C-D). Interestingly, genetically defined ICN types are not evenly distributed throughout the heart, with a spatial preference for *Ddah1*^+^ ICNs being more enriched in the posterior ganglia (Figure 1E-F). Our data also revealed a range of receptors, ion channels, and neurochemicals in the ICNS (Figure S1B), some of which are differentially expressed in *Ddah1*^+^ and *Npy*^+^ ICNs, suggesting distinct roles for these populations in cardiac regulation. To validate our single-cell data, we further performed RNAscope HiPlex analysis. As expected, *Ddah1* and *Npy* were expressed in largely non-overlapping ICNs, with similar percentages to those observed in single-cell RNA sequencing (Figure 1G). *Ddah1* was often co-expressed with *Dbh*, while *Npy* co-expressed with *Ptger3*. *Vip* was only expressed in 2.0 ± 0.4 % ICNs, mainly co-expressing with *Npy*. Also consistent with single-cell RNA-sequencing data, *Ddah1*^+^ ICNs were more abundant in the posterior regions, with the highest concentration observed in one specific ganglion (Figures 1G and S2).

To gain genetic access to various ICN subtypes, we generated Ddah1-2a-Cre^ERT2^ and Dbh-2a-Cre^ERT2^ mice, and acquired Npy-ires-Cre^42^, Vip-ires-Cre^43^, and Slc18a2-Cre^44^ mouse lines (Figures 2A, S3). Consistent with our single cell RNA sequencing results, 30.0 % ICNs were labelled in Ddah1^tdT^ mice, and these co-expressed *Ddah1* (98.7 %) but not *Npy* (0.6 %), as confirmed by RNAscope (Figure 2B). Strong fluorescence was observed in brain regions including olfactory areas and the thalamus in Ddah1^Ai203^ mice (Figure S3C). Similarly, most ICNs labelled in Dbh^Ai203^ mice co-expressed *Ddah1* (72.7 %) but not *Npy* (Figure S3D). As expected, adrenergic neurons in the locus coeruleus, nucleus tractus solitarii (NTS), area postrema (AP), and rostral ventrolateral medulla (RVLM) were labelled in Dbh^tdT^ mice (Figure S3E), and sympathetic postganglionic neurons in the stellate ganglia were extensively labelled in both Dbh^tdT^ and Npy^tdT^ mice (Figure S3F-G). In the hearts of Npy^tdT^ and Npy^Ai203^ mice, reporters predominantly labeled ICNs that co-expressed *Npy* (Figures 2B, S3H). Only a few ICNs were labelled in Vip^L10-GFP^ and Slc18a2^sun1-GFP^ mice (Figure 2D-E). These findings indicate that our mouse models accurately target the corresponding ICN populations. Collectively, our data reveal two primary genetically distinct ICN types and provide a means to access their genetic characteristics.

**Figure 2.**
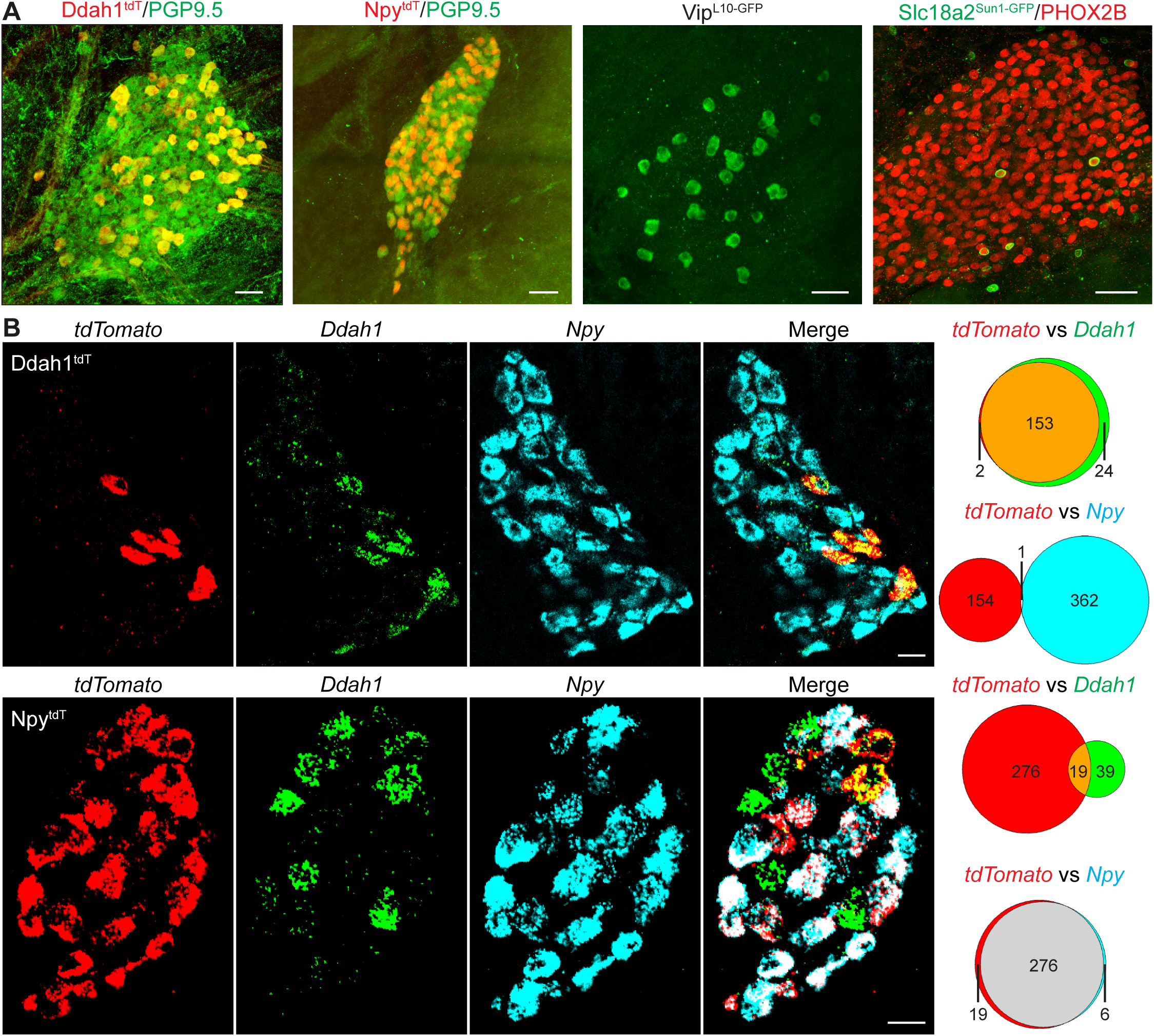
Mouse genetic toolkits for accessing various ICN subtypes. (A) ICN ganglia from Ddah1^tdT^, Npy^tdT^, Vip^L10-GFP^, and Slc18a2^Sun1-GFP^ mice, showing reporter-labeled ICNs co-stained with pan-ICN markers PGP9.5 or PHOX2B. (B), RNAscope Multiplex Assays performed on ICN ganglia from Ddah1^tdT^ (top) and Npy^tdT^ (bottom) mice, using probes against *tdTomato* (red), *Ddah1* (green), and *Npy* (cyan). The numbers of cells expressing each gene alone or in combination were quantified. Scale bars, 50 μm (A), 20 μm (B). See also Figure S3.

### Npy^+^ and Ddah1^+^ ICNs have distinct projections patterns

To investigate whether the genetic segregation of *Ddah1*^+^ and *Npy*^+^ ICNs reflects functional differences, we compared their single-cell transcriptomic profiles with annotated neuron populations from the well-characterized ENS. We integrated our ICN single cell RNA-seq dataset with a published EN dataset^41^ (Figure S4). EN subtypes were defined by canonical markers, including *Tac1* (excitatory motor neurons), *Nos1*/*Npy*/*Vip* (inhibitory motor neurons), *Nmu*/*Cck*/*Nxph2* (intrinsic primary afferent neurons, or IPANs), and *Sst*/*Neurod6*/*Dbh* (interneurons). Notably, most *Npy*^+^ ICNs clustered with EN inhibitory motor neurons, suggesting that they may function as motor neurons, whereas *Ddah1*^+^ ICNs showed transcriptomic similarity to multiple EN sensory neuron and interneuron types. This cross-organ analysis supports the idea that *Npy*^+^ and *Ddah1*^+^ ICNs are functional distinct.

To further characterize these subtypes, we examined their anatomical projection patterns within the heart. We developed a novel cell type-specific anatomical tracing strategy using adeno-associated virus (AAV) to selectively label ICN subtypes (Figure 3A). As a proof of principle, we injected AAV9-flex-tdTomato into the heart’s atrial fat pad of Baf53b-Cre mice via transverse sternotomy^45^ (referred to as Baf53b^AAV-tdT^). This method effectively labelled ICNs and their projections (Figures 3A and S5A-B), enabling high-resolution mapping of neuronal morphology and innervation patterns. To compare projection profiles of *Npy*^+^ and *Ddah1*^+^ ICNs, we performed analogous AAV injections in Npy-ires-Cre and Ddah1-2a-Cre^ERT2^ mice to generate Npy^AAV-tdT^ and Ddah1^AAV-tdT^ mice respectively. We then used iDISCO-based tissue clearing^46^ followed by lightsheet microscopy to visualize and quantify their projection patterns in whole-mount hearts (Figure 3B). As expected, neuronal cell bodies were effectively labelled in both lines (Figure S5C). *Npy*^+^ ICNs exhibited broad innervation throughout the heart, with dense projections to the atrial appendages, upper and lower ventricles extending to the apex, and around the pulmonary veins (Figure 3C). Notably, they robustly innervated both the sinoatrial (SA) and atrioventricular (AV) nodes, suggesting roles in regulating cardiac rhythm and conduction. In addition, *Npy*^+^ ICNs formed prominent fiber networks around the aortic root, indicating potential involvement in regulating outflow tract dynamics. In contrast, *Ddah1*^+^ ICNs showed a markedly more restricted innervation pattern, with dense fibers only observed in a few specific domains: the left atrium, the pulmonary veins, and the pulmonary trunk. Unlike the widespread and uniform pulmonary vein innervation seen in *Npy*^+^ ICNs, *Ddah1*^+^ ICNs primarily targeted the pulmonary vein ostia (Figures 3C, S5D), a site commonly associated with arrhythmogenesis and atrial fibrillation^47^, suggesting a potential role in electrophysiological modulation. *Ddah1*^+^ ICNs also showed dense innervation of the pulmonary trunk, a region minimally innervated by *Npy*^+^ neurons. Interestingly, although *Ddah1*^+^ ICNs densely innervated the AV node, they provided little innervation to the SA node, suggesting that they may influence cardiac rhythm and conduction through mechanisms distinct from *Npy*^+^ ICNs. Quantitative analysis revealed that compared to *Npy*^+^ ICNs, *Ddah1*^+^ ICNs exhibited significantly lower projection density per neuron, defined as total projection signal normalized by the number of labeled neurons, in vast majority of heart regions (Figure 3D), indicating more spatially restricted communication with cardiac cells. In contrast, *Ddah1*^+^ ICNs displayed denser fiber connectivity across ICN ganglia (Figure 3E-F), pointing to a potential role as interganglionic interneurons specialized for local circuit integration. Together, these anatomical tracing studies uncover striking divergence in the cardiac projection architecture of *Npy*^+^ and *Ddah1*^+^ ICNs, highlighting distinct functional specializations within the ICNS.

**Figure 3.**
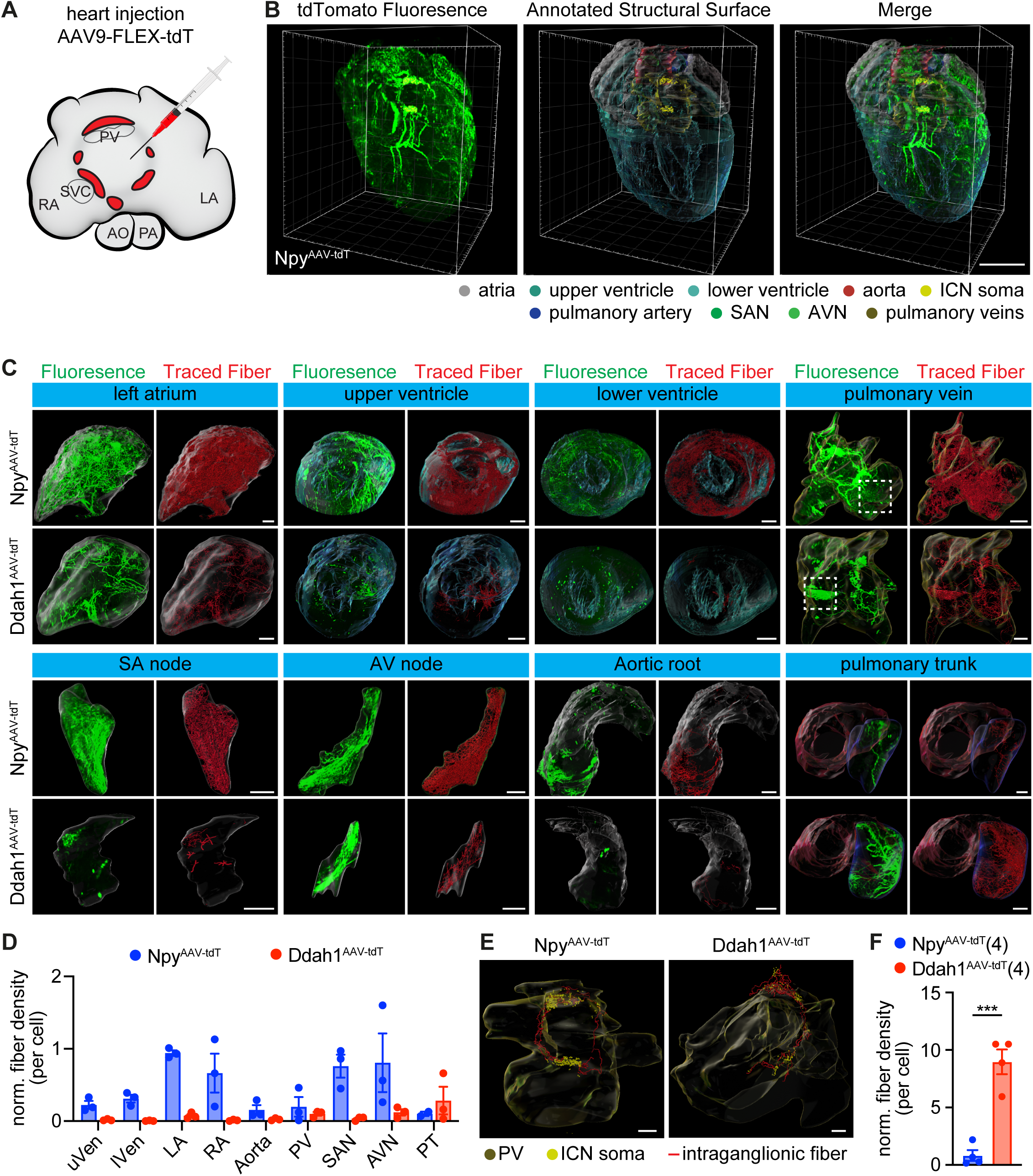
*Npy*⁺ and *Ddah1*⁺ ICNs Exhibit Distinct Cardiac Projection Patterns. (A) Schematic of cell type-specific anatomical tracing of genetically defined ICN subtypes via intracardiac AAV injection. (B) Lightsheet imaging of a cleared Npy^AAV-tdT^ heart showing tdTomato-labeled projections (green) overlaid with annotated cardiac structures (color-coded as indicated). (C) Representative images from Npy^AAV-tdT^ and Ddah1^AAV-tdT^ hearts showing region-specific innervation patterns. Green, tdTomato fluorescence; red, segmented nerve fibers. Zoomed-in regions (dashed boxes) are shown in Figure S5D. (D) Quantification of normalized fiber density per neuron across indicated cardiac regions in Npy^AAV-tdT^ and Ddah1^AAV-tdT^ mice. (E) Representative images of labeled ICN soma (yellow dots) and intraganglionic nerve fibers (red) in Npy^AAV-tdT^ and Ddah1^AAV-tdT^ hearts. (F) Quantification of normalized intraganglionic fiber density per neuron. Data represent mean ± SEM. Significance: ***p<0.001, two-tailed Student’s t test. Scale bars: 2 mm (B), 0.5 mm (C, D). See also Figures S4 and S5.

### Npy^+^ but not Ddah1^+^ ICNs are parasympathetic postganglionic neurons

While it is well established that cardiac parasympathetic postganglionic neurons are part of the ICNS, their precise identities remain unclear. To pinpoint the ICNs involved in parasympathetic functions, we examined *Fos* expression using RNAscope in Baf53b^L10-GFP^ mice in response to the baroreflex induced by intraperitoneal (i.p.) injection of phenylephrine (PE)^6,8,9^, which potently reduced heart rate via activation of parasympathetic postganglionic ICNs (Figure 4A-F). In control mice treated with saline, *Fos* expression was minimal, observed in only 0.2 % of ICNs, consistent with previous findings indicating low baseline parasympathetic activity in mice^6^. In contrast, in mice treated with phenylephrine (PE), *Fos* was highly expressed in 54.9 ± 2.4 % ICNs, with the vast majority (93.9 ± 1.8 %) of these *Fos*^+^ ICNs co-expressing *Npy* but not *Ddah1* (5.2 ± 2.0 %). From a different perspective, most (71.5 ± 5.1 %) of *Npy^+^* ICNs while 12.3 ± 4.0 % of *Ddah1*^+^ ICNs responded to PE. These results indicate that *Npy*^+^ ICNs, but not *Ddah1*^+^ ICNs, are involved in cardiac parasympathetic regulation.

**Figure 4.**
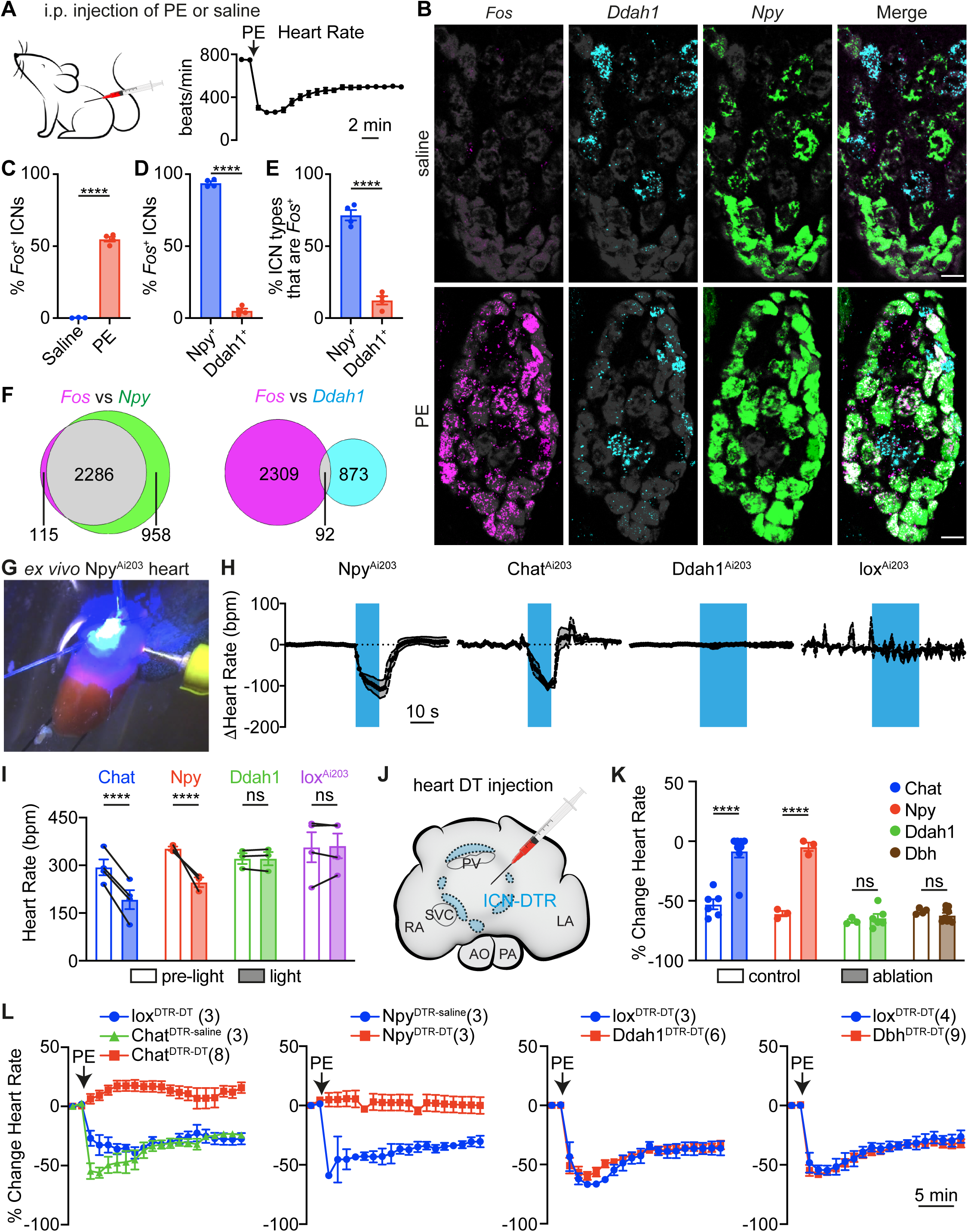
*Npy*^+^ but not *Ddah1*^+^ ICNs are cardiac postganglionic parasympathetic neurons. (A) Schematic of PE-induced baroreflex (left), illustrating the prolonged decrease in heart rate (right). Data shown as mean ± SEM, from 8 mice. (B) RNAscope HiPlex Assays on ICN ganglia from Baf53b^L10-GFP^ mice treated with saline (top) or PE (bottom), using probes against *Fos* (magenta), *Ddah1* (cyan), and *Npy* (green). GFP-labeled ICNs are shown in grey, marking the full ICN population. Scale bars, 20 μm. (C-F) Quantification of RNAscope experiments as in (B): (C) Percentage of *Fos*^+^ ICNs in saline- and PE-treated mice. (D) Percentage of *Fos*^+^ ICNs that are *Npy*^+^ or *Ddah1*^+^ in PE-treated mice. (E) Percentage of *Npy*^+^ and *Ddah1*^+^ ICNs that are *Fos*^+^. (F) Numbers of cells expressing each gene alone or in combination. Data represent mean ± SEM, from 3 saline- and 4 PE-treated mice. (G) Representative image of *ex vivo* optogenetic stimulation of ICNs on a retrogradely perfused Langendorff heart from Npy^Ai203^ mice. (H) Heart rate responses to optogenetic activation (blue bar) of ICN subtypes. (I) Quantification of heart rate changes from (H). Data shown as mean ± SEM, from 4 Chat^Ai203^, 3 Npy^Ai203^, 3 Ddah1^Ai203^, and 4 lox^Ai203^ mice. (J) Schematic illustration of ICN ablation via local injection of DT into the atrial fat pad of appropriate DTR-expressing mice via transverse sternotomy. (K, L) Quantification (K) and temporal dynamics (L) of PE-induced heart rate changes (expressed as a percentage) after ICN subtype ablation. Data as mean ± SEM, from indicated animal numbers. Statistics: two-tailed Student’s t test (C-E), two-way ANOVA followed by Šidák’s multiple comparisons test with paired (I) or non-paired (K) comparisons. Significance: ns, not significant; *p<0.05; **p<0.01; ***p<0.001; ****p<0.0001. See also Figure S6.

To confirm the parasympathetic function of *Npy*^+^ ICNs, we then tested whether optogenetic stimulation of these neurons could reduce heart rate. To eliminate confounding influences from extrinsic autonomic inputs, we took advantage of the Ai203 mice^48^, which express a Cre-dependent soma-targeted opsin sChroME, and performed optogenetic experiments in an *ex vivo* retrogradely perfused Langendorff heart setup (Figure 4G). Notably, optogenetic activation of both *Chat*^+^ and *Npy*^+^ ICNs in isolated hearts from Chat^Ai203^ and Npy^Ai203^ mice rapidly and significantly reduced heart rate, whereas no heart rate changes were observed in Ddah1^Ai203^ and Cre-negative controls upon light stimulation (Figure 4H-I). These findings demonstrate that *Npy*^+^, but not *Ddah1*^+^, ICNs regulate heart rate, consistent with our anatomical data showing that only *Npy*^+^ ICNs innervate the SA node (Figure 3C). Interestingly, although both *Npy*^+^ and *Ddah1*^+^ ICNs innervate the AV node, only optogenetic activation of *Npy*^+^ ICNs resulted in a prolonged PR interval (Figure S6A-C). This further confirms that *Ddah1*^+^ ICNs lack parasympathetic function and likely influence cardiac electrophysiology via non-parasympathetic mechanisms.

We next investigated whether *Npy*^+^ ICNs are required for parasympathetic control of the heart. To selectively ablate specific ICN subtypes, we employed diphtheria toxin (DT)-mediated ablation using iDTR mice, which carry a Cre-dependent DT receptor (DTR) allele^49^. In this well-established system, only Cre-expressing cells express DTR and are selectively eliminated upon DT administration, allowing precise and targeted cell ablation^9,49^. To restrict ablation to the ICNS, we locally administered DT into the atrial fat pad using the same surgical approached previously employed for viral delivery (Figure 4J), thereby ensuring regional specificity. Ablation of *Chat*^+^ ICNs in Chat^DTR^ mice (Chat^DTR-DT^) effectively abolished the PE-induced bradycardia, confirming successful disruption of parasympathetic output, while no effect was observed in saline-treated (Chat^DTR-saline^) or Cre-negative (lox^DTR-DT^) littermate control mice (Figures 4K-L, S6D). Notably, ablation of *Npy*^+^ ICNs in Npy^DTR-DT^ mice also exhibited a complete loss of PE-induced bradycardia, whereas Npy^DTR-saline^, Ddah1^DTR-DT^, Dbh^DTR-DT^, and their respective littermate control mice showed normal responses (Figures 4K-L, S6E-G). These results demonstrate that *Npy*^+^ ICNs, but not *Ddah1*^+^/*Dbh*^+^ ICNs, are essential for parasympathetic regulation of heart rate. Together, these findings identify *Npy*^+^ ICNs as the functional relevant cardiac parasympathetic postganglionic neurons in the ICNS, distinguishing them from *Ddah1*^+^ ICNs which appear to serve non-parasympathetic roles.

### *Npy^+^* ICNs are essential for cardiac stability and survival

Unexpectedly, Chat^DTR-DT^ and Npy^DTR-DT^ mice, which lack parasympathetic ICNs, died within one week of DT administration. In contrast, no adverse effects were observed in Ddah1^DTR-DT^, Dbh^DTR-DT^, and multiple control groups including lox^DTR-DT^, Chat^DTR-saline^, and Npy^DTR-saline^ mice (Figure 5A). Importantly, Chat^DTR^ mice that received systemic DT via i.p. injection at the same dose did not show similar lethality (Figure 5A), demonstrating that the observed effect is due to local ICN ablation rather than systemic toxicity. In both Chat^DTR-DT^ and Npy^DTR-DT^ mice, heart rate began to decline ∼48 hours post-DT, followed progressive reductions in respiration rate and body temperature, despite stable blood oxygen saturation (Figures 5B, S7A). Death consistently occurred when heart rate fell to ∼300 beats per minute (bpm) and body temperature approached ambient levels. In contrast, Ddah1^DTR-DT^, Dbh^DTR-DT^, and control mice maintained stable physiological parameters throughout. This paradoxical heart rate decline was not due to excessive acetylcholine release during neuronal death, as atropine treatment, which effectively blocks baroreflex-induced bradycardia (Figure S7B), failed to prevent the heart rate drop or improve survival (Figure S7C-D).

**Figure 5.**
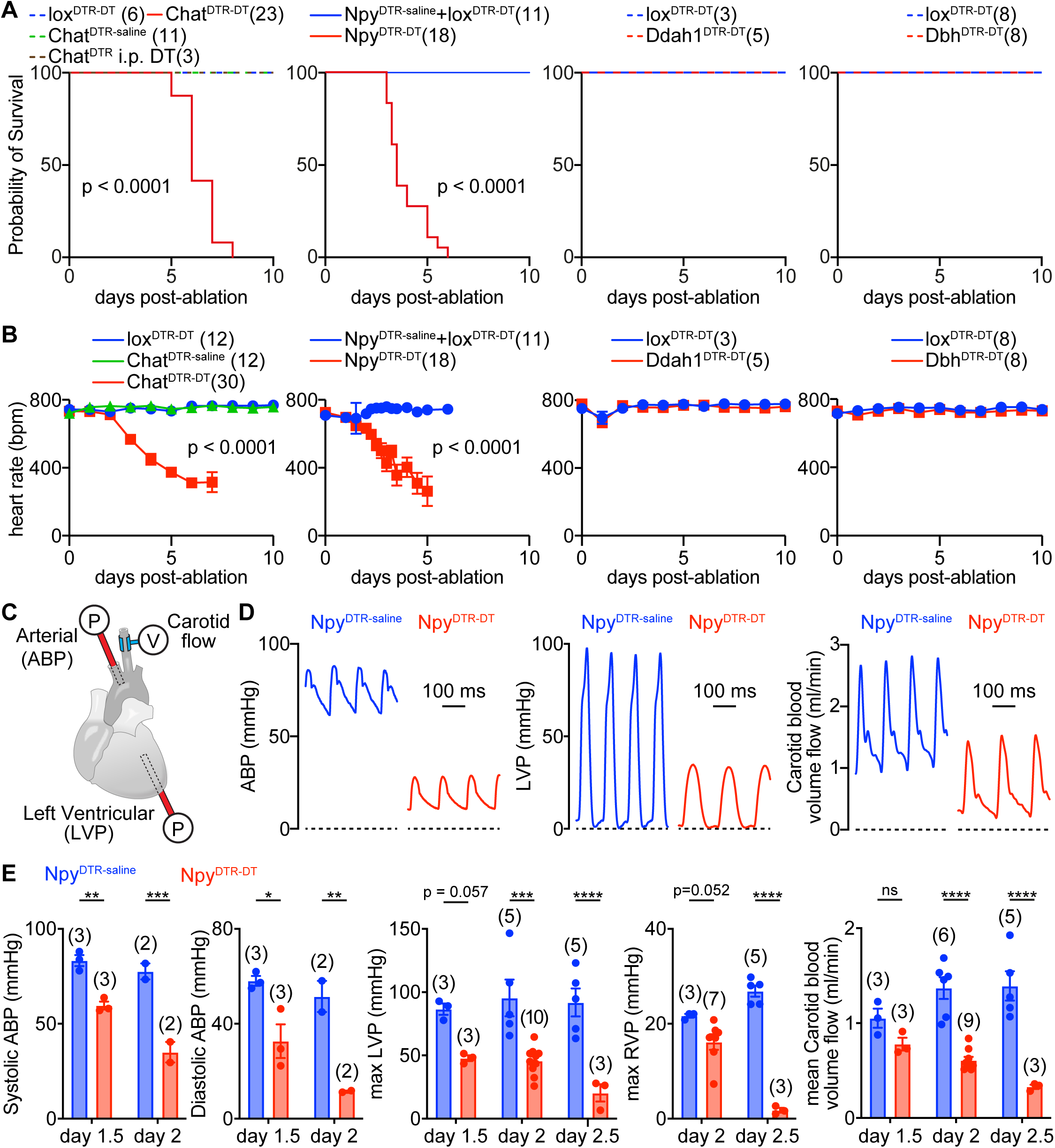
*Npy*⁺ ICNs sustain cardiac stability and support survival. (A) Survival curve of animals following ablation of specific ICN subtypes. Experimental conditions and number of animals are indicated. (B) Heart rate changes in animals after ICN subtype ablation. Data shown as mean ± SEM, from indicated number of animals. (C) Schematic illustration of measurements for arterial blood pressure (ABP), left ventricular pressure (LVP), and carotid blood flow. (D) Representative traces of ABP, LVP, and carotid blood volume flow in Npy^DTR-saline^ and Npy^DTR-DT^ mice, 48 hours post-injection. (E) Quantification of cardiovascular parameters in Npy^DTR-saline^ and Npy^DTR-DT^ mice at the indicated time points post-injection. Data presented as mean ± SEM, from the indicated number of animals. Statistics: log-rank (Mantel-Cox) test (A), one-way ANOVA (B), two-way ANOVA followed by Šidák’s non-paired multiple comparisons test (E). Significance: *p<0.05; **p<0.01; ***p<0.001; ****p<0.0001. See also Figure S7.

To investigate the mechanisms underlying the progressive heart rate decline, we closely monitored cardiac function at multiple time points following *Npy*^+^ ICN ablation (Figures 5C-E, S7E-F). Notably, by 36 hours post DT administration in Npy^DTR-DT^ mice, prior to any detectable heart rate reduction (Figure 5B), significant decreases in arterial blood pressure (ABP) and left ventricular pressure (LVP) were already evident. Specifically, systolic ABP decreased by 28.4 ± 2.5%, diastolic ABP by 43.8 ± 12.3%, and LVP by 44.8 ± 2.7%, compared with littermate Npy^DTR-saline^ controls, suggesting an early impairment in cardiac contractility. In contrast, carotid arterial blood flow and integrated blood flow per heartbeat, a proxy for stroke volume, remained largely unchanged (Figures 5E, S7E-F).

Cardiac function continued to deteriorate over the next 24 hours. By 48 hours post-DT, systolic ABP in Npy^DTR-DT^ mice had dropped 54.8 ± 3.3 %, and diastolic ABP declined 77.7 ± 0.3 % relative to Npy^DTR-saline^ control mice (Figure 5D-E). By 60 hours post-DT, carotid arterial blood flow had decreased by 76.4 ± 1.6 %, LVP by 77.9 ± 7.4 %, and right ventricular pressure (RVP) by 93.4 ± 1.8 %. Despite the profound cardiac dysfunction, heart rhythm and PR interval remained stable (Figure S7G-I), indicating that the conduction system was intact. Similar physiological deterioration was observed in Chat^DTR-DT^ mice, but not in their Chat^DTR-saline^ littermate controls (Figure S7J-L). Together, these findings demonstrate that disruption of *Npy*^+^ ICNs results in severe, life-threatening cardiac dysfunction, underscoring their essential role in maintaining normal cardiac performance and autonomic homeostasis.

### ICNs maintain aortic root mechanics to support coronary perfusion

To better understand the cellular mechanism underlying cardiac dysfunction triggered by ICN ablation, we next measured cardiac function in *ex vivo* Langendorff heart preparations from ICN-ablated and control mice (Figure 6A-B). Surprisingly, while *in vivo* cardiac function was markedly impaired in Chat^DTR-DT^ mice (Figures 5B, S7J-L), these deficits were not observed *ex vivo*, where hearts were retrogradely perfused with sufficient nutrients and oxygen. Specifically, heart rate, LVP, RVP, and coronary vascular resistance, represented as mean perfusion flow under constant perfusion pressure, were comparable between Chat^DTR-DT^ mice and their littermate controls (Figures 6B). These results suggest that ICN ablation does not impair myocardial contractility, SA node frequency, or cardiac conduction when adequate perfusion is maintained.

**Figure 6.**
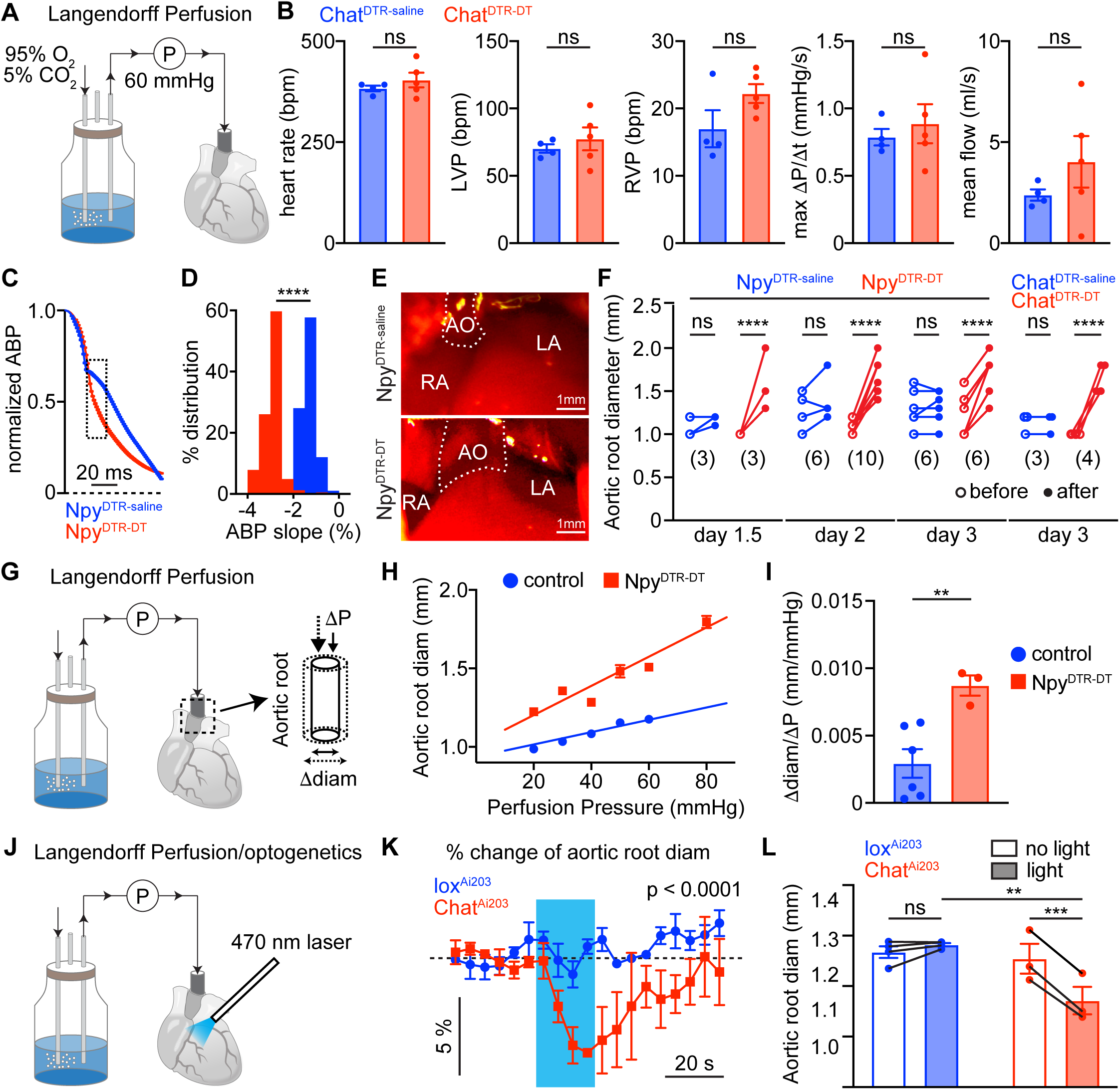
ICNs regulate aortic root dynamics. (A) Schematic illustration of the *ex vivo* retrogradely perfused Langendorff heart preparation. (B) Quantification of cardiac parameters from Langendorff-perfused hearts of Chat^DTR-saline^ and Chat^DTR-DT^ mice, demonstrating preserved cardiac functions with adequate perfusion. Data shown as mean ± SEM, from indicated number of animals. (C) Normalized arterial blood pressure (ABP) traces from Npy^DTR-saline^ and Npy^DTR-DT^ mice. Data as mean ± SEM, representing 200 cardiac cycles per animal, from 3 mice per group. (D) Distribution of ABP slope values from regions highlighted by the dashed rectangle in (C), illustrating altered pressure dynamics in the absence of *Npy*^+^ ICNs. (E) Representative images of the aortic root in Npy^DTR-saline^ and Npy^DTR-DT^ mice. Dashed lines outline the aortic root (AO); LA, left atrium; RA, right atrium. Scale bars, 1 mm. (F) Quantification of aortic root diameter before and at indicated time points following ICN subtype ablation. Sample sizes are noted. (G) Schematic illustration of the experimental setup to assess the relationship between perfusion pressure and aortic root diameter in *ex vivo* Langendorff hearts. (H) Graph showing perfusion pressure vs aortic root diameter for representative Npy^DTR-DT^ (red) and control (blue) mice. For each mouse, diameters were measured across multiple pressure points. A linear regression line was fitted to calculate the slope, representing mechanical compliance of the aortic root. (I) Quantification of aortic root compliance, based on slope measurements. Data shown as mean ± SEM, from 6 control and 3 Npy^DTR-DT^ mice. (J) Schematic illustration of *ex vivo* optogenetic stimulation of ICNs. (K) Temporal dynamics of percentage change in aortic root diameter in response to optogenetic stimulation (blue bar) in Chat^Ai203^ and control mice. (L) Quantification of optogenetic responses, as shown in (K). Data represent mean ± SEM, from 4 control and 3 Chat^DTR-DT^ mice. Statistics: two-tailed Student’s t test (B, D, I, K), two-way ANOVA followed by Šidák’s paired multiple comparisons test (F, L). Significance: ns, not significant; **p<0.01; ***p<0.001; ****p<0.0001.

Since myocardial perfusion primarily occurs during diastole rather than systole^50,51^, it depends heavily on coronary resistance and aortic diastolic pressure. In particular, the rise in aortic pressure after aortic valve closure, visible as the dicrotic notch, helps sustain the pressure gradient between the aorta and the coronary ostia located just above the aortic valve, thereby promoting coronary flow during early diastole^52^. While coronary resistance remained unchanged (Figure 6B), we observed a marked reduction in both magnitude (Figure 5E) and kinetics (Figure 6C-D) of diastolic aortic pressure in Npy^DTR-DT^ mice. Specifically, the dicrotic notch was absent, and the pressure declined more rapidly after aortic valve closure, both of which would be expected to impair coronary perfusion. To understand the mechanistic basis of this altered pressure profile, we examined the geometry and compliance of the aortic root, which plays a critical role in buffering pressure during diastole. Notably, Npy^DTR-DT^ and Chat^DTR-DT^ mice showed significant aortic root enlargement (Figure 6E-F), suggesting a loss of elastic recoil needed to sustain diastolic pressure. To directly assess mechanical changes of the aortic root, we further examined its pressure-volume relationship in *ex vivo* Langendorff heart preparations, where perfusion pressure could be precisely controlled (Figure 6G). Indeed, aortic root compliance, measured as Δdiameter/Δpressure, was significantly increased in Npy^DTR-DT^ hearts compared to controls (Figure 6H-I), confirming that the aortic root had become more distensible, resembling an easily expandable balloon. Together, these findings demonstrate that ICNs are essential for preserving the mechanical properties of the aortic root that maintain diastolic pressure and enable effective coronary perfusion.

These functional results align with our anatomical finding that *Npy*^+^ ICNs tightly encircle the aortic root, a feature not seen in Ddah1^AAV-tdT^ mice (Figure 3C), raising the possibility that ICNs actively regulate aortic root mechanical tone in real time. To test this, we performed optogenetics in *ex vivo* Langendorff hearts from Chat^Ai203^ mice while imaging the aortic root at high spatial resolution using a high-speed camera (Figure 6J). Strikingly, optogenetic stimulation of ICNs for 20 seconds led to a gradual reduction in aortic root diameter, an effect absent in their lox^Ai203^ littermate controls (Figure 6K-L), indicating that ICNs can dynamically modulate aortic root mechanics, potentially adjusting cardiac outflow dynamics to promote coronary perfusion during parasympathetic responses. Together, these findings reveal that ICNs play an active and critical role in tuning aortic root dynamics to ensure adequate coronary perfusion and support overall cardiac function, highlighting their broader importance in cardiovascular homeostasis.

### Ddah1^+^ ICNs preserve cardiac resilience during extreme stress

To uncover the physiological role of *Ddah1*^+^ ICNs, we next examined functional impairments in Ddah1^DTR-DT^ mice, in which these neurons are selectively ablated. Although these mice maintained normal heart rate, respiration, and general activity at baseline, they displayed striking vulnerability under stress. In a physical restraint test, a well-established model of psychological stress in rodents^53^, animals were confined in a tube (2.5 cm in diameter) for 20 minutes daily over 10 days. While all control lox^DTR-DT^ mice (14/14) survived without abnormal behavior, most Ddah1^DTR-DT^ mice (12/19, 63.2%) died within six days (Figure 7A). Notably, all deaths occurred during the restraint period and within minutes of onset, suggesting sudden cardiac failure. To further investigate the underlying cardiac events, we built a specialized restraint chamber allowing continuous ECG recording in conscious, restrained mice (Figure S8A). ECG traces revealed that affected animals showed a rapid drop in heart rate beginning 1-3 minutes before death (Figure 7C-D), accompanied by arrhythmias, first- and second-degree AV block, and cardiac irregularities (Figure S8B). These findings confirm that Ddah1^DTR-DT^ mice died from sudden cardiac arrest during stress induced by physical restraint.

**Figure 7.**
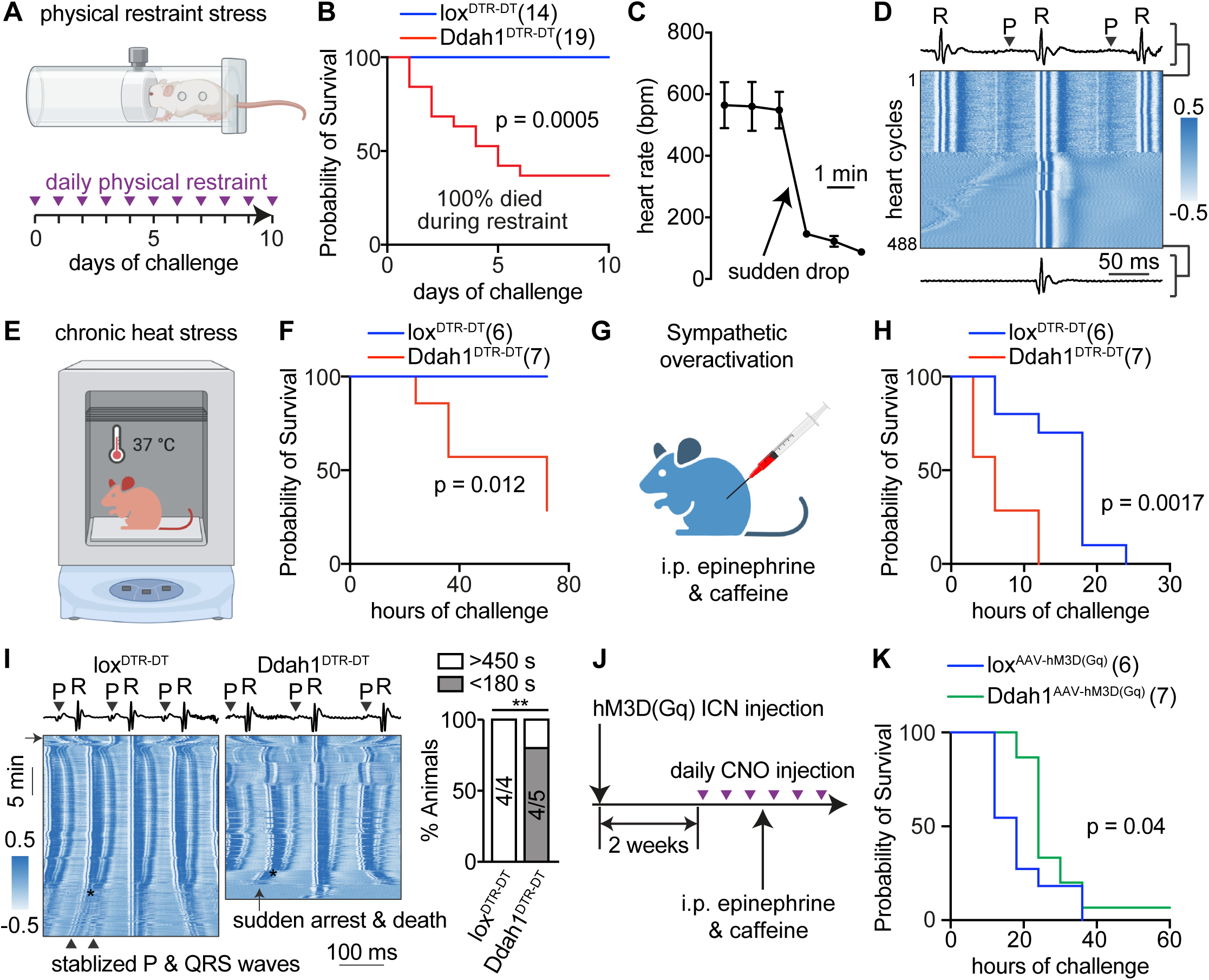
*Ddah1*^+^ ICNs are essential for protecting the heart during extreme stress. (A) Schematic illustration of the physical restraint test. Mice were restrained daily for 20 minutes for up to 10 days. (B) Survival curve of mice subjected to physical restraint. Sample numbers are indicated. (C) Heart rate dynamics prior to death in *Ddah1*^+^ ICN ablated mice. Data as mean ± SEM, from 3 Ddah1^DTR-DT^ mice. (D) Representative heatmap showing ECG signal from a Ddah1^DTR-DT^ mouse recorded shortly before death. Each row represents a 300 ms ECG segment centered on the R wave of a detected heart cycle. Time is aligned to the R wave peak to visualize temporal patterns and evolving ECG dynamics across cycles. Signal amplitude is color-coded to reflect waveform morphology. ECG traces corresponding to the top and bottom rows of the heatmap are displayed above and below the plot, respectively, with P and R waves labeled to indicate their position and color representation in the heatmap. (E) Schematic of the chronic heat stress test. (F) Survival curve of mice challenged with chronic heat stress. Sample numbers are indicated. (G) Schematic of the sympathetic overactivation test. (H) Survival curve of mice challenged with sympathetic overactivation. Sample numbers are indicated. (I) Representative heatmaps showing ECG signals from lox^DTR-DT^ (control) and Ddah1^DTR-DT^ mice during the sympathetic overactivation test. ECG trace corresponding to the top row of the heatmap is shown above, with P and R waves labeled. Arrow on the left indicates the time of epinephrine & caffeine injection, which triggered a rapid reduction in RR interval. While 4/4 control mice survived >450 s after AV dissociation (marked by *), 4/5 Ddah1^DTR-DT^ mice died within 180 s, demonstrating severe cardiac instability. (J) Schematic of the chronic *Ddah1*^+^ ICN stimulation protocol. (K) Survival curve showing that chronic stimulation of *Ddah1*^+^ ICNs significantly improves survival under sympathetic overactivation. Sample numbers are indicated. Statistics: log-rank (Mantel-Cox) test (B, F, H, K), Chi-square test (I). Significance: **p<0.01. See also Figure S8.

We then challenged Ddah1^DTR-DT^ mice with additional types of stressors. Epidemiological studies have demonstrated that chronic heat exposure significantly increases the risk of cardiovascular diseases, including heart attack and sudden death^54^. We therefore housed mice at 37°C over a 7-day period (Figure 7E). While all control lox^DTR-DT^ mice (6/6) survived this mild but sustained thermal stress, most Ddah1^DTR-DT^ mice (5/7, 71.4%) died within the first three days (Figure 7F). To test whether Ddah1^DTR-DT^ mice exhibited heightened susceptibility to sympathetic overactivation, a known trigger of arrhythmia and sudden cardiac death^55,56^, we administered a lethal dose of epinephrine and caffeine (100 mg/kg and 120 mg/kg, respectively)^57^. Remarkably, Ddah1^DTR-DT^ mice exhibited significantly accelerated mortality compared to control lox^DTR-DT^ littermates (Figure 7G-H). To assess cardiac electrical changes, we repeated the epinephrine/caffeine challenge in anesthetized mice and continuously measured their ECG. Both groups showed shortened R-R intervals consistent with increased heart rate, as well as arrythmia and gradual P-R interval prolongation, which eventually leads to AV dissociation (Figure 7I). While most Ddah1^DTR-DT^ mice (4/5) experienced sudden cardiac arrest within three minutes following AV dissociation, all control mice (4/4) were able to temporarily stabilize cardiac rhythms and reestablish AV coupling, surviving for longer periods. Together, these results demonstrate that *Ddah1*^+^ ICNs are essential for maintaining cardiac rhythm and electrical stability in response to diverse stress challenges.

We next asked whether enhancing *Ddah1*^+^ ICN activity could improve survival during extreme stress. To test this, we used a chemogenetic strategy involving Designer Receptors Exclusively Activated by Designer Drugs (DREADDs). We selectively expressed the excitatory hM3D(Gq) receptor in *Ddah1*^+^ ICNs by injecting AAV9-Flex-hM3D(Gq)-mCherry into the hearts of Ddah1-2a-Cre-ERT2 mice (Ddah1^AAV-hM3D(Gq)^), following the same approach described above. As a proof of principle, robust mCherry expression in Chat^AAV-hM3D(Gq)^ mice confirmed effective targeting of ICNs two weeks after viral delivery, and these neurons could be reliably activated by clozapine N-oxide (CNO), a widely used DREADD ligand (Figure S8C). To test for protective effects under stress, Ddah1^AAV-hM3D(Gq)^ mice received daily CNO (i.p.) beginning two weeks post-injection and were then challenged with the same lethal dose of epinephrine/caffeine three days into CNO treatment (Figure 7J). Strikingly, Ddah1^AAV-hM3D(Gq)^ mice showed significantly enhanced survival compared to their lox^AAV-hM3D(Gq)^ littermate controls (Figure 7K), demonstrating that chronic activation of *Ddah1*^+^ ICNs can confer robust protection against extreme cardiac stress. Together, these findings establish *Ddah1*^+^ ICNs as critical mediators of cardiac resilience and identify them as promising therapeutic targets for preventing stress-induced sudden cardiac death.

## Discussion

Neurocardiac interactions are essential for maintaining optimal heart function across a range of physiological conditions^1,2^. The ICNS, a central component of the heart-brain axis, plays a vital role in regulating autonomic balance and is increasingly recognized as a contributor to cardiac pathologies such as arrhythmias and heart failure^2,12,24^. In this study, we identify two molecularly distinct neuronal subtypes within the ICNS, each indispensable for survival yet specialized for different aspects of cardiac control, one sustaining day-to-day autonomic homeostasis, the other ensuring resilience during stress. Together, our findings highlight the biological and clinical importance of the ICNS as both a fundamental regulator of cardiac function and a promising target for therapeutic intervention.

Many visceral organs, including the gut, heart, lung, and pancreas, contain intrinsic nervous systems that were historically viewed as simple relay stations for parasympathetic input from the vagus nerve. However, recent studies have redefined this view, particularly in the ENS, where extensive molecular, anatomical, and functional diversity has revealed a complex, semi-autonomous neural network often referred to as the “second brain” of the gut^58^. Inspired by these findings, and by observations that ICNs exhibit neurochemical and functional heterogeneity, it has been speculated that the ICNS may similarly comprise sensory, inter-, and motor neurons forming a functionally independent network or “little brain” on the heart. Our results support this idea in part, revealing that while the ICNS is clearly more than a simple parasympathetic relay, its molecular diversity appears more limited than that of the ENS^41,59,60^. For example, *Npy*^+^ ICNs resemble ENS inhibitory motor neurons, and *Ddah1*^+^ ICNs share transcriptional features with certain sensory or interneuron-like populations in the ENS. In contrast, the ICNS lacks excitatory motor neurons and the broad spectrum of sensory and interneuron subsets found in the gut. Moreover, many canonical sensory receptors and ion channels (e.g., *Piezo1*, *Piezo2*, *Trpv1*, *Calca*) are largely absent from ICNs, suggesting that the ICNS may rely on distinct or unconventional sensory mechanisms. These differences highlight the organ-specific adaptations of intrinsic nervous systems, reflecting the distinct physiological demands and regulatory priorities of each host organ.

While all visceral organs require coordinated regulation, the heart has unique physiological demands due to its critical role in pumping blood continuously throughout life, requiring precise, reliable, and adaptable control. Our study reveals that two molecularly distinct populations of ICNs are specialized to meet different physiological demands: *Npy*^+^ ICNs provide stable, beat-to-beat regulation under basal conditions, whereas *Ddah1*^+^ ICNs support adaptive cardiac responses during extreme or heightened physiological states. Notably, these populations exhibit distinct anatomical organizations (Figure 3). *Npy*^+^ ICNs display broader anatomical distribution and significantly larger average innervation fields compared to *Ddah1*^+^ ICNs, suggesting a structural basis for their divergent functional roles. Generally, coordinated regulation of organ function can be achieved through one of two strategies: distinct neurons control specific functions and coordinate through extensive local interactions, enabling flexibility but reducing stability; or individual neurons regulate multiple functions across broader territories, enhancing stability at the cost of specificity. While some degree of spatial specificity in the ICNS has been previously reported^15,17,61,62^, the expansive innervation fields of *Npy*⁺ ICNs suggest a model in which individual neurons contribute to the regulation of multiple cardiac functions across wide regions of the heart. This architecture may help explain how the ICNS achieves robust control despite its limited neuronal complexity, particularly in contrast to systems like the ENS, which relies on a far more intricate network of interneurons. Together, these findings provide a framework for understanding how different organizational strategies within organ-intrinsic neural networks can support both stable and flexible control of vital organ function.

The ICNS is increasingly recognized for its essential role in cardiac medicine and has been implicated in the pathogenesis of several major cardiac disorders, including atrial fibrillation, heart failure, and sudden cardiac death. For example, the pulmonary vein region, densely innervated by ICNs, is a well-established hotspot for arrhythmogenesis, and catheter ablation of this area is a common clinical intervention for atrial fibrillation^21,63,64^. However, a major limitation in both experimental and therapeutic approaches is the lack of cell-type specific tools, with most studies relying on non-selective electrical stimulation or broad radiofrequency or cryoablation. This lack of precision has contributed to confounding and sometimes contradictory results. Consistent with these limitations, clinical trials of vagus nerve stimulation for heart failure and sudden cardiac death have produced largely disappointing outcomes^65–68^. Our discovery of Ddah1⁺ ICNs introduces a new, previously unrecognized mechanism for adaptive regulation under stress or pathophysiological conditions, independent of classical parasympathetic pathways. Although the exact mechanisms remain to be determined, these neurons likely function either by acting downstream of sympathetic overdrive to provide negative feedback that restrains excessive cardiac excitation, or by serving as intrinsic sensory neurons that detect perturbations in cardiac performance and initiate localized corrective responses. Nevertheless, the identification of *Ddah1*⁺ ICNs opens a promising new avenue for advancing cardiac medicine. It not only expands our understanding of neural mechanisms underlying heart protection but also offers a framework for identifying novel cellular therapeutic targets. Ultimately, this discovery lays the foundation for developing more selective, mechanism-based interventions for heart disease, with the potential to overcome the limitations of current non-specific autonomic therapies.

## Resource Availability

For further information and requests, please contact the lead contact, Rui Chang. Ddah1-2a-Cre^ERT2^ and Dbh-2a-Cre^ERT2^ mice will be deposited at the Jackson Laboratory and made available upon request. Single cell RNA-seq data generated for the current study will be made publicly available at NCBI Gene Expression Omnibus (GEO). Code for Seurat will be available at github.com.

## Acknowledgements

We thank D. Spergel, C. Yu, and Q. Zhao for valuable insights on the project, L. Ziolkowski, Y. Nikolaev, and S. Bagriantsev for optogenetic testing of soma-targeted opsins, S. Wilson, L. Shao, and Yale CNNR Imaging Core for assistance with microscopy, M. Schneeberger for assistance with lightsheet imaging analysis, G. Wang and YCGA for assistance with single-cell sequencing, J. Greenwood, P. Shamble, and Yale Neuroscience Technology Core for making imaging apparatus and chambers. Funding was provided by NIH (DP2HL151354, R01HL150449 to R.B.C, R01HL148008 to L.H.Y., DP2DA056169 to L. Z., T32GM136651, F30AT013179 to O.A.H.), the G. Harold & Leila Y. Mathers Foundation (R.B.C.), the Allen Discovery Center program, a Paul G. Allen Family Foundation (R.B.C. and L.Z.), Cancer Prevention and Research Institute of Texas grant (RR200084 to X.Z.), Yale School of Medicine Science Fellowship (P.E.RC), and Kavli Postdoctoral Fellowship (I.H.).

## Author Contribution

Q.J.X. and R.B.C. designed experiments, analyzed data, and wrote the manuscript. Q.J.X. led and performed all experiments. M.C.A. designed and built ECG-compatible restraint chamber. O.A.H. contributed to RNAscope experiments and analysis. I.H. performed immunohistochemistry for Slc18a2^Sun1-GFP^ mice. R.P.K performed brain cryosectioning. P.E.RC contributed to lightsheet imaging analysis. R.L.W. contributed to surgeries and immunostaining. Q.J.X., I.H., L.Z., and R.B.C. analyzed single cell RNA-seq data. Q.J.X., M.C.A., and R.B.C. performed all other data analysis. L.H.Y., X.Z., and R.B.C. supervised heart functional studies.

## Declaration of interests

The authors declare no competing interests.

## Declaration of generative AI and AI-assisted technologies

The authors used ChatGPT4o to check for grammar.

## Supplemental information

**Figure S1.**
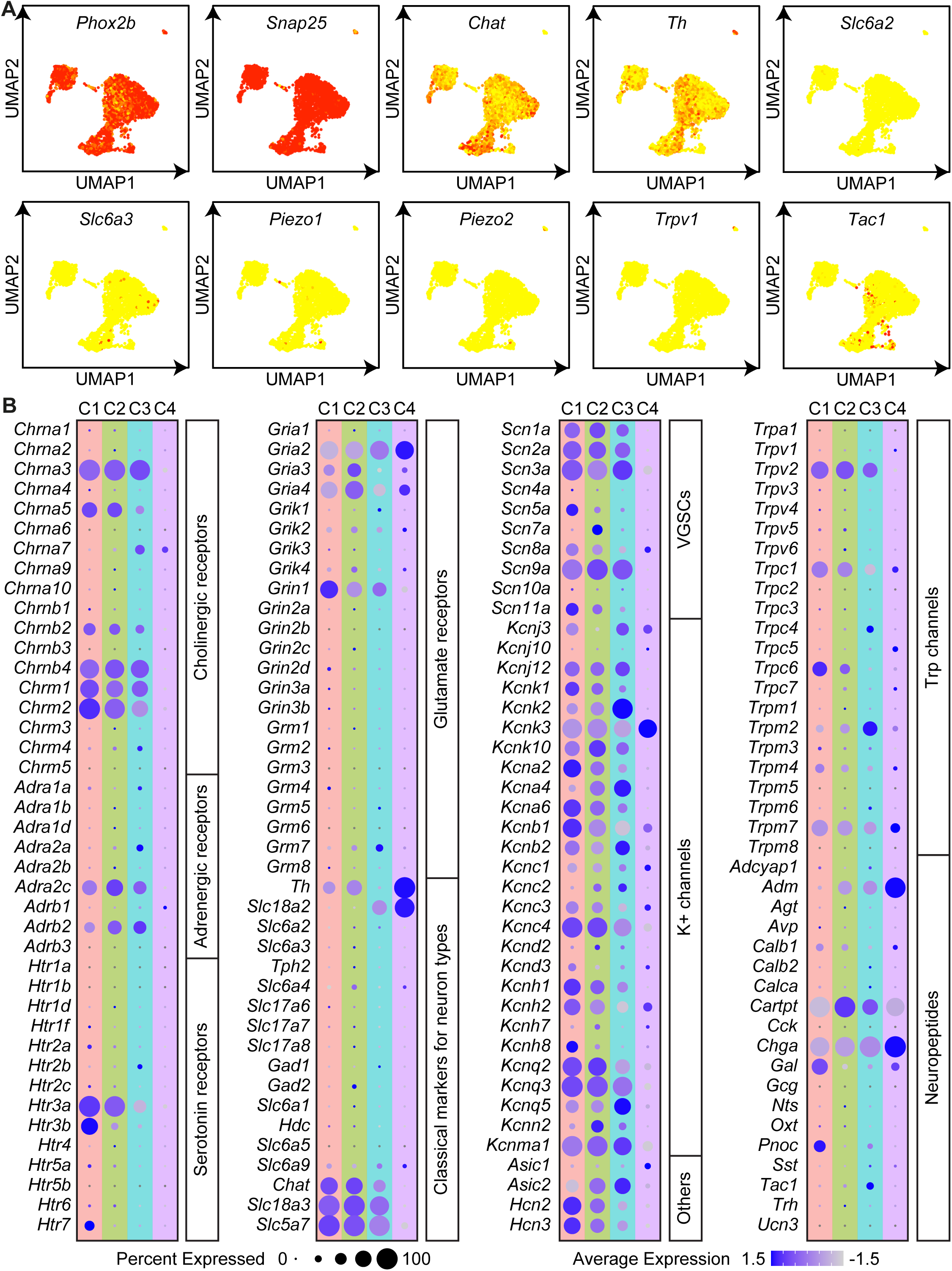
Expression of selected receptors, ion channels, and neuropeptides in the ICNS, related to Figure 1. (A) UMAP plots, as shown in Figure 1B, colored by the expression of the indicated marker genes. (B) Dot plots showing the expression of selected receptors, ion channels, neuropeptides, and other signaling molecules across ICN subtypes.

**Figure S2.**
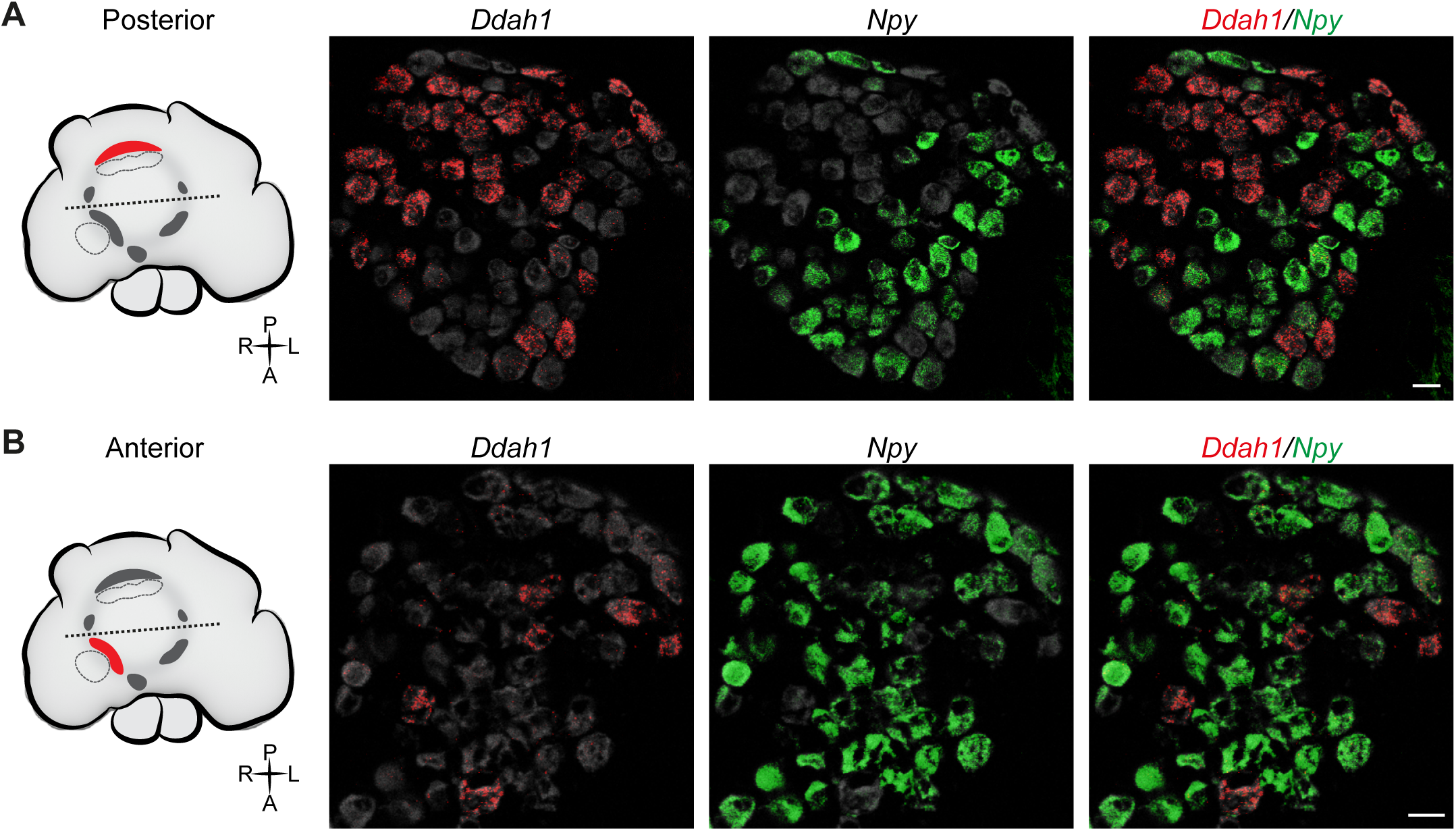
Spatial segregation of ICN subtypes in cardiac ganglia, related to Figure 1. (A-B) RNAscope Multiplex Assays using probes against *Ddah1* (red) and *Npy* (green) on posterior (A) and anterior (B) ICN ganglia from Baf53b^L10-GFP^ mice, as illustrated in the accompanying schematic. GFP-labeled ICNs are shown in grey, marking the full ICN population. Images show spatial enrichment of distinct ICN subtypes across ganglia. Scale bars, 20 μm.

**Figure S3.**
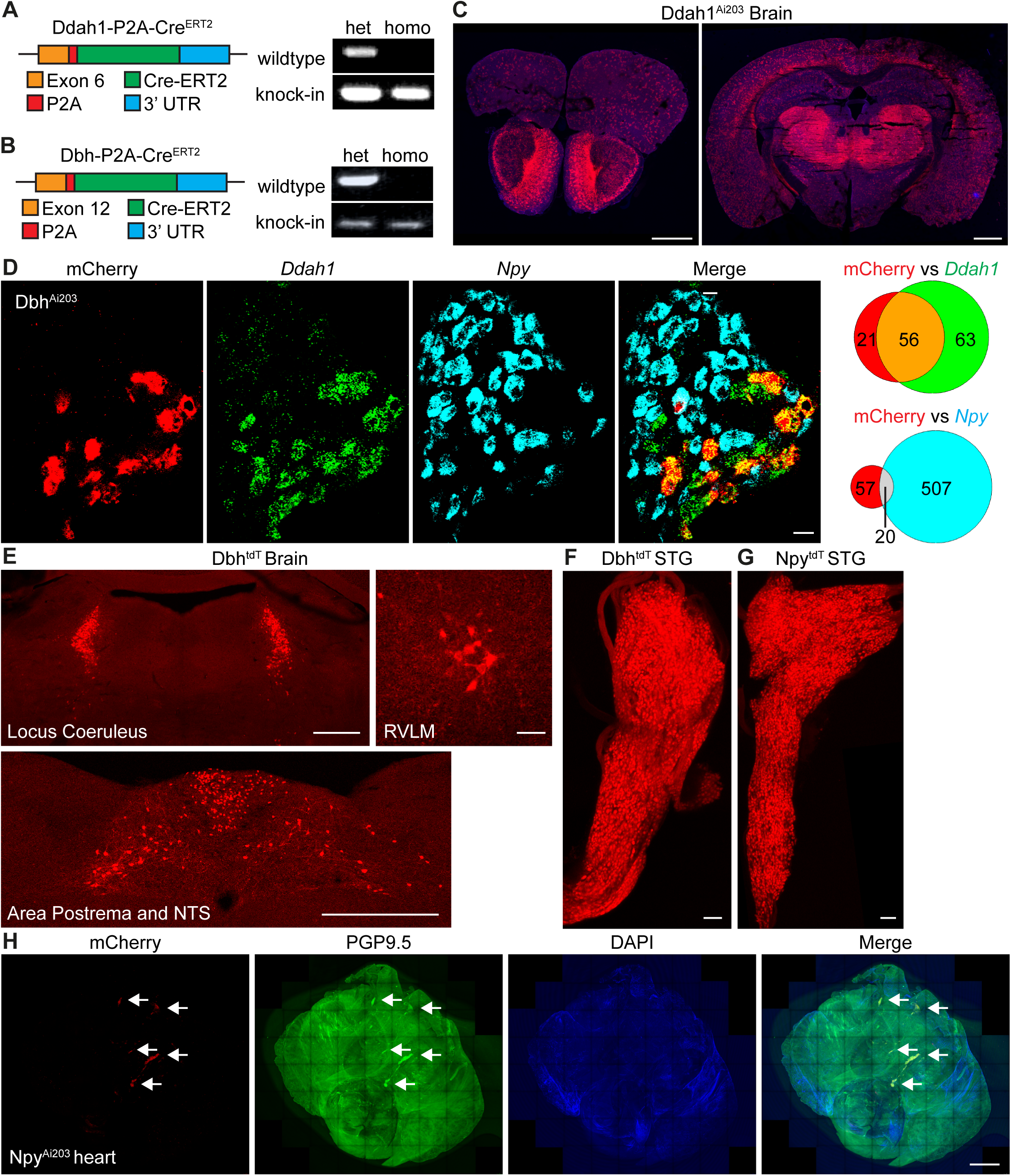
Characterization of Cre mouse lines for targeting ICN subtypes, related to Figure 2. (A-B) Schematic design of Ddah1-2a-Cre^ERT2^ (A) and Dbh-2a-Cre^ERT2^ (B) mouse lines. Genomic PCR with allele-specific primers confirmed the presence of wild-type and knock-in alleles in each line. (C) Representative brain cryosections from Ddah1^Ai203^ mice showing mCherry expression. (D) RNAscope Multiplex Assay on ICN ganglia from Dbh^Ai203^ mice using probes for *Ddah1* (green) and *Npy* (cyan) and immunostaining for mCherry (red). Cells expressing each gene individually or in combination were quantified. (E) tdTomato expression in selected brain regions of Dbh^tdT^ mice, including the locus coeruleus, rostral ventrolateral medulla (RVLM), nucleus tractus solitarii (NTS), and area postrema. (F-G) Stellate ganglia from Dbh^tdT^ and Npy^tdT^ mice, showing robust tdTomato expression in sympathetic postganglionic neurons. (H) Wholemount image of a flattened heart atrium from Npy^Ai203^ mice, showing mCherry expression localized to PGP9.5^+^ ICN ganglia. Scale bars: 1 mm (C, H), 20 μm (D), 100 μm (E, F), 50 μm (G, RVLM), 500 μm (G, locus coeruleus and area postrema).

**Figure S4.**
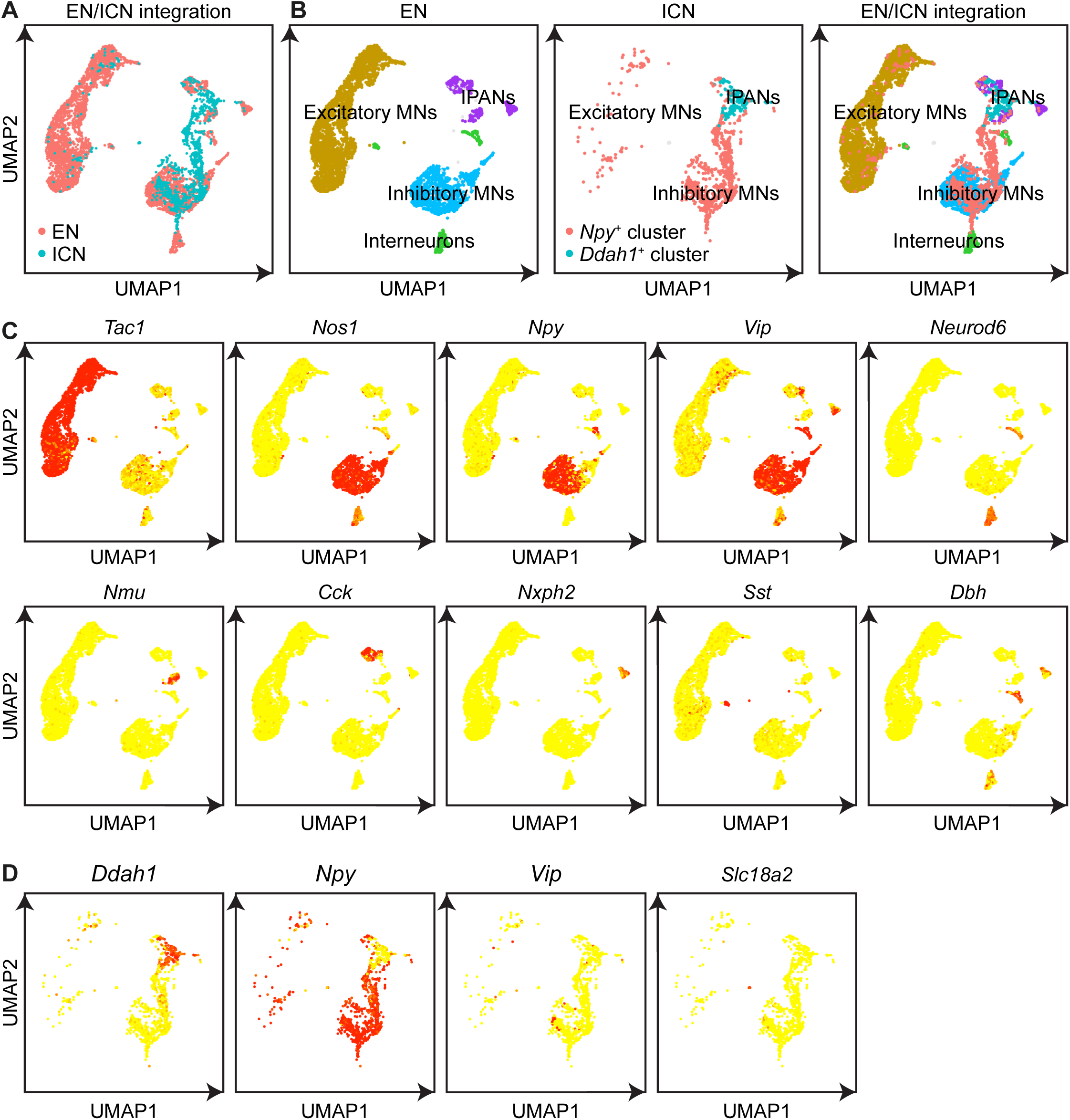
Cross-organ single cell integration highlights functional features of ICN subtypes, related to Figure 3. (A) UMAP plot showing integrated single cell RNA-seq datasets of 1,111 ICNs and 4,316 enteric neurons (ENs), colored by sample origin. (B) UMAP plots displaying ENs only, ICNs only, and the integrated dataset, colored by annotated EN subtypes and *Ddah1*^+^ or *Npy*^+^ ICN subtypes. *Npy*^+^ ICNs colocalize with EN inhibitory motor neurons, while *Ddah1*^+^ ICNs colocalize with intrinsic primary afferent neurons (IPANs). (C) Expression patterns of selected marker genes used to define EN subtypes. (D) Expression of key marker genes distinguishing ICN subtypes.

**Figure S5.**
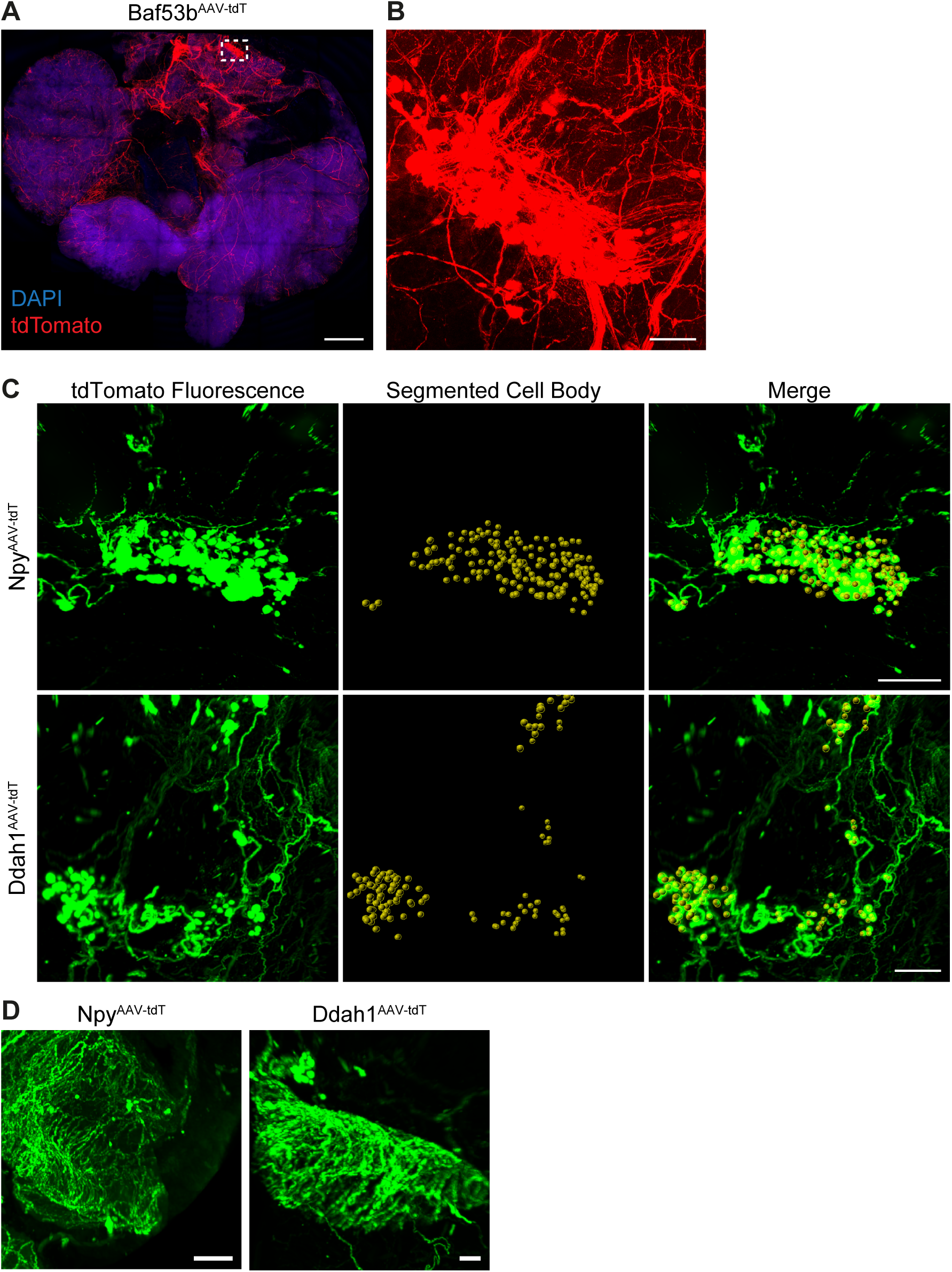
Cell type-specific labeling of *Npy*⁺ and *Ddah1*⁺ ICN, related to Figure 3. (A) Flattened whole-mount Baf53b^AAV-tdT^ heart showing robust labeling of ICN soma and nerve fibers. (B) Higher magnification of the dashed region in (A). (C) tdTomato-labeled ICN soma (green) and their segmented positions (yellow dots) in Npy^AAV-tdT^ and Ddah1^AAV-tdT^ hearts. (D) Zoomed-in images of the boxed regions from Figure 3C, highlighting regional innervation patterns. Scale bars: 1 mm (A), 200 μm (B, D), 500 μm (C).

**Figure S6.**
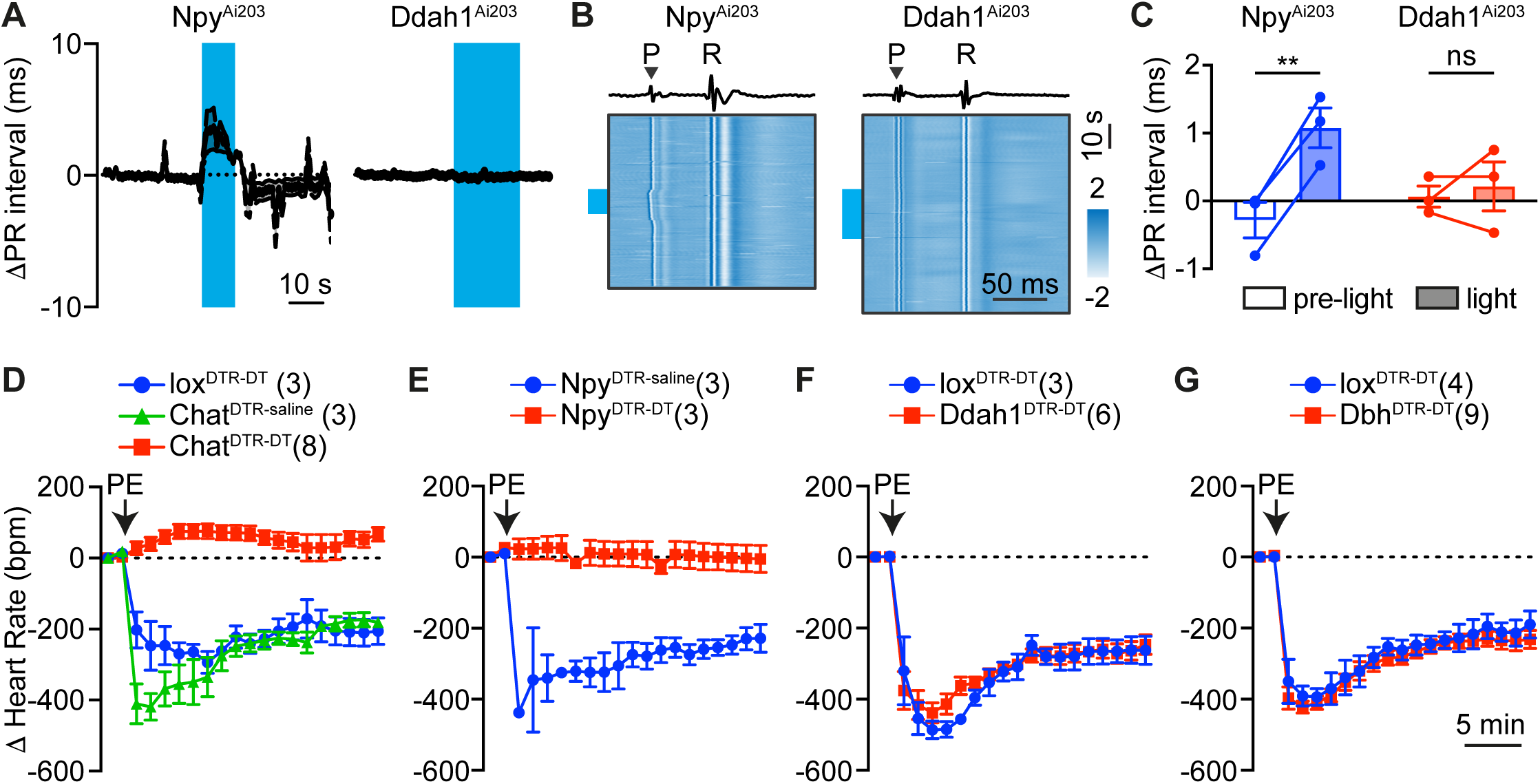
Manipulation of ICN subtypes confirms parasympathetic identity of *Npy*^+^ ICNs, related to Figure 4. (A) Temporal dynamics of PR interval responses to optogenetic activation (blue bar) of ICN subtypes. Data as mean ± SEM, n = 3 mice per group. (B) Representative heatmaps showing ECG signals from Npy^Ai203^ and Ddah1^Ai203^ mice during optogenetic stimulation (indicated by blue bars at left). Each row represents a 200 ms ECG segment centered on the R wave of a detected heart cycle. Time is aligned to the R wave peak to visualize temporal patterns and evolving ECG dynamics across cycles. Signal amplitude is color-coded to reflect waveform morphology. The ECG trace corresponding to the top row of the heatmap is shown above, with P and R waves labeled. (C) Quantification of PR interval changes before and during light stimulation of ICN subtypes. Data as mean ± SEM, n = 3 mice per group. (D) Temporal dynamics of PE-induced heart rate changes (in beats per minute) following ICN subtype ablation. Data as mean ± SEM. Sample sizes are indicated. Statistics: two-way ANOVA followed by Šidák’s paired multiple comparisons test (C). Significance: ns, not significant; **p<0.01.

**Figure S7.**
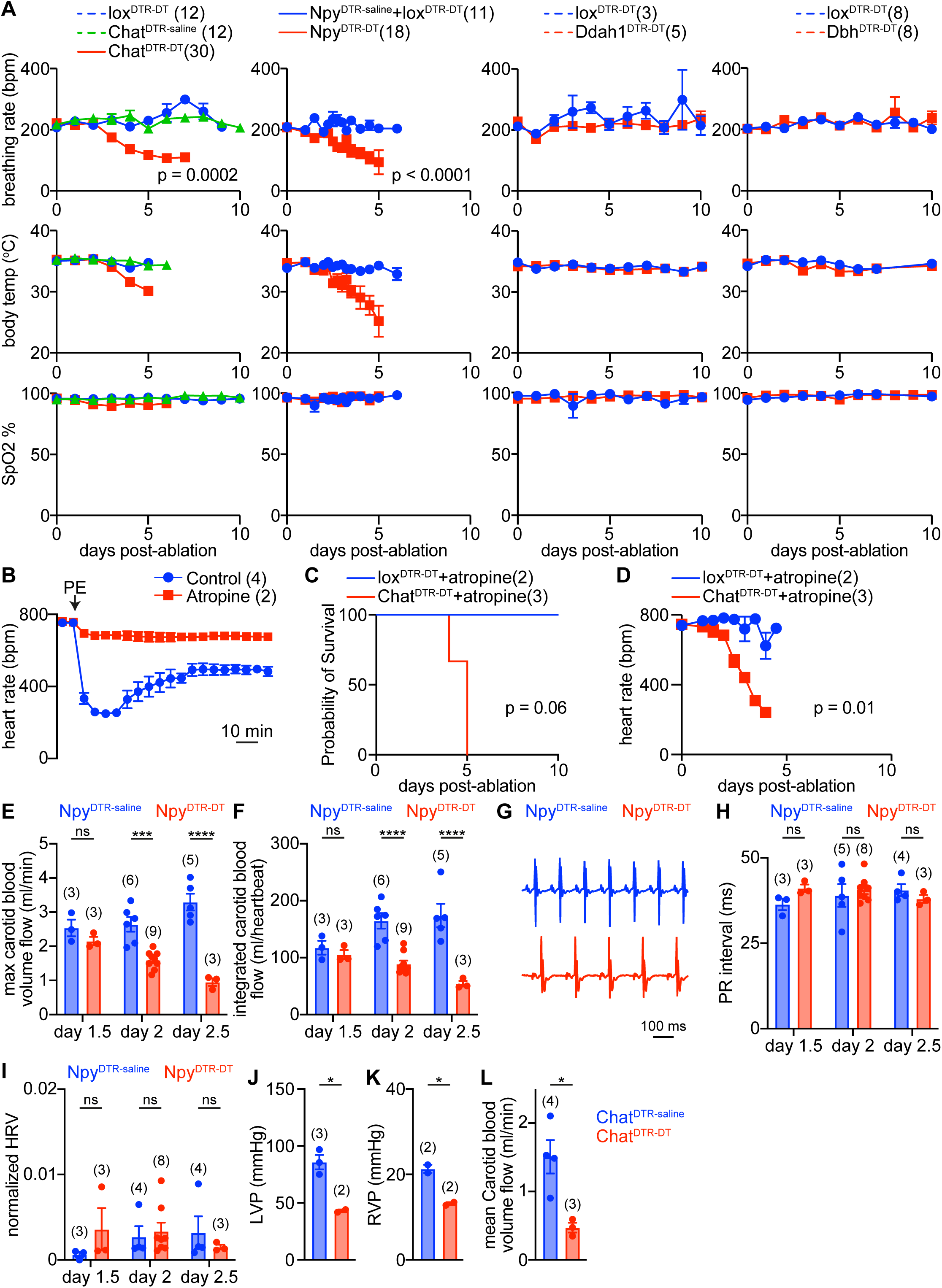
Roles of *Npy*^+^ ICNs in cardiac homeostasis and survival, related to Figure 5. (A) Physiological changes in animals after ICN subtype ablation. (B) Temporal dynamics of heart rate during PE stimulation, showing that PE-induced bradycardia is abolished by atropine treatment. (C) Survival curve of atropine-treated animals after pan-ICN ablation. Sample sizes are indicated. (D) Heart rate changes in atropine-treated animals following pan-ICN ablation. (E-F) Quantification of maximum carotid blood volume flow (E) and integrated carotid blood flow per heartbeat (F) in Npy^DTR-saline^ and Npy^DTR-DT^ mice at indicated time points post-injection. (G) Representative ECG traces from Npy^DTR-saline^ and Npy^DTR-DT^ mice. (H) Quantification of PR interval at indicated time points post-injection in Npy^DTR-saline^ and Npy^DTR-DT^ mice. (I) Quantification of normalized heart rate variability (HRV) in Npy^DTR-saline^ and Npy^DTR-DT^ mice at indicated time points post-injection. (J-L) Quantification of cardiovascular parameters including left ventricular pressure (LVP; J), right ventricular pressure (RVP; K), and mean carotid blood flow (L) in Chat^DTR-saline^ and Chat^DTR-DT^ mice at 72 hours post-injection. All quantitative data are presented as mean ± SEM, with sample sizes indicated for each analysis. Statistic: one-way ANOVA (A, Chat), two-tailed Student’s t test (A, Npy; D, J-L), log-rank (Mantel-Cox) test (C), two-way ANOVA followed by Šidák’s non-paired multiple comparisons test (E, F, H, I). Significance: ns, not significant; *p<0.05; ***p<0.001; ****p<0.0001.

**Figure S8.**
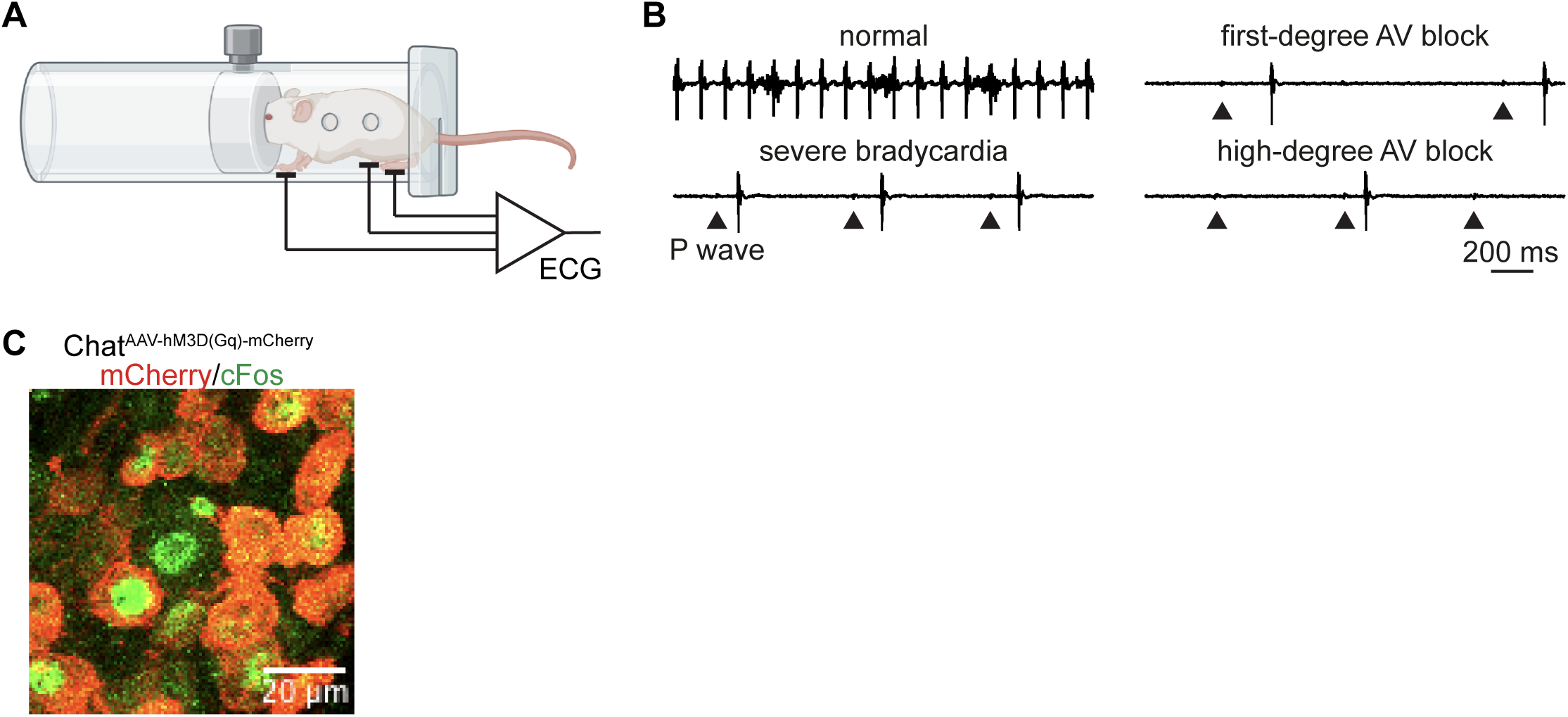
Assessment of *Ddah1*^+^ ICN function during stress, related to Figure 7. (A) Schematic of a custom-designed restraint chamber for ECG recording in restrained, awake mice. (B) Representative ECG traces showing normal rhythm and abnormal events including various degrees of AV blocks observed immediately prior to death. Arrow heads indicate P waves. (C) ICN ganglia from Chat^AAV-hM3D(Gq)-mCherry^ mice, showing selective induction of cFos (green) in infected, mCherry^+^ ICNs (red) 30 minutes after chemogenetic stimulation. Scale bar: 20 μm.

## Methods

### Animals

All animal husbandry and experimental procedures were conducted in compliance with the Yale University’s Institutional Animal Care and Use Committee (IACUC) and National Institute of Health (NIH) guidelines. All efforts were made to minimize animal pain and reduce the number of animals used. Mice were housed under standard 12:12 hour light-dark cycles with ad libitum access to food and water. All animals used were age- and sex-matched adults (older than 8 weeks), and no sex-specific differences were observed.

#### Mouse lines

The following mouse lines were obtained from the Jackson Laboratory: wildtype C57BL/6J (000664), Npy-ires-Cre (027851), Baf53b-Cre (027826), Chat-ires-Cre (031661), Vip-ires-Cre (010908), lox-tdTomato (007914), lox-iDTR (007900), Ai203 (037939), Sun1-GFP (021039). Slc18a2-Cre mice were obtained from MMRRC (037427). lox-L10-GFP mice were described before^34,35,69^. Ddah1-2a-Cre^ERT2^ and Dbh-2a-Cre^ERT2^ mouse lines were custom-generated by Taconic Biosciences.

#### Tamoxifen treatment

To induced reporter expression in Ddah1^DTR^, Ddah1^Ai203^, Ddah1^tdT^, Dbh^tdT^ and Dbh^DTR^ mice, animals were gavaged with tamoxifen (Toronto Chemical research) dissolved in corn oil at a dose of 100 mg/kg once daily for five consecutive days starting at 4 weeks of age. For anatomical and neuronal manipulation studies, Ddah1^AAV-tdT^ and Ddah1^AAV-hM3D(Gq)^ mice were gavaged with tamoxifen (100 mg/kg in corn oil) beginning one day prior to heart injection surgeries and continuing for five consecutive days.

### Single-cell RNA sequencing and data analysis

#### ICN isolation and sequencing

ICNs were acutely isolated using a modified protocol for peripheral neurons^32,69^. Briefly, hearts were harvested from 10 age- and sex-matched Baf53b^GFP^ mice into ice-cold Leibovitz’s L-15 Medium (Gibco^TM^). ICN-containing regions were identified and dissected under a Leica M205FCA Fluorescence Stereo Microscope. Tissues were trimmed into 1-3 mm^2^ pieces and incubated with 0.25% trypsin-EDTA (Gibco^TM^) at 37 °C for 25 minutes. After washing in L-15 medium supplemented with 10% fatal bovine serum (FBS), tissues were digested with 1 mg/mL Collagenase A (Sigma) and 3 mg/mL of Dispase II (Sigma) in 1 x Hank’s Balanced Salt Solution (HBSS, Stemcell Technologies) at 37°C for 40 minutes. Dissociated tissues were triturated using a series of three fire-polished Pasteur pipettes of decreasing opening size, then filtered through a 40 μm cell strainer (Corning). Cells were centrifuged at 200 x g for 10 minutes at 4°C and resuspended in 1x HBSS containing 0.04% BSA. For regional transcriptomic analysis shown in Figure 1E, anterior and posterior ganglia were dissected and processed separately. For mixed ganglia samples, tissues were pooled. Fluorescently labelled cells were sorted using a BD FACS Aria cell sorter into ice-cold 1x HBSS with 0.04% BSA at the Yale Flow Cytometry Facility. Approximately 5,000-10,000 sorted cells were loaded per channel on the 10x Genomics Chromium microfluidic device to target a recovery of 3,000-6,000 cells per sample. Single-cell cDNA libraries were prepared using the Chromium Single Cell 3’ V3 reagent kit at the Yale Center for Genomic Analysis (YCGA) and sequenced on an Illumina NovaSeq S4 sequencer at 150-300 million reads to achieve a target depth of 30,000-50,000 reads per cell.

#### Bioinformatic processing

Raw sequencing data were aligned to the mm10-2020-A mouse genome reference using the Cell Ranger software v.7.1.0 (10X Genomics). Cells were filtered using the following quality control thresholds: number of detected genes per cell > 2,500 and < 20,000; and percentage of mitochondria genes < 10%. To select neuronal populations, cells with *Plp1* expression ≥1 and Fabp7 ≥1 were excluded, and those expressing Phox2b >0 and Snap25 >0 were retained. A total of 2,870 high-quality ICNs passed filtering (1,111 from mixed ganglia, 933 from anterior ganglia, and 826 from posterior ganglia). Filtered datasets were integrated and analyzed using the Seurat R package v.4.4.2^36^. Dimensionality reduction was performed using the top 50 principal components, and UMAP^37^ was used for visualization. ICN cell types (e.g., in Figure 1B) were annotated based on unsupervised clustering. DEGs between ICN subtypes were identified using the Seurat’s built-in likelihood-ratio test based on the bimod distribution. For comparative analysis with enteric neurons (Figure S4), single-cell RNA-sequencing data from a previously published study^41^ were reprocessed using the same quality control and filtering pipeline. A total of 4,316 enteric neurons and 1,111 ICNs (from mixed ganglia) were integrated and analyzed together. Enteric neuron subtypes were annotated based on previously defined marker genes, while ICN clusters were identified using cell barcodes corresponding to subtype annotations from Figure 1B.

### RNAscope Assay

RNAscope Hiplex Assays (Figures 1G, 4B, and S2) were performed using the Hiplex Fluorescent Assay v2 kit (Advanced Cell Diagnostics) following the manufacturer’s protocol, as previously described^32,69^. Cardiac regions containing ICNs were carefully dissected from Baf53b^GFP^ animals using a Leica M205FCA Fluorescence Stereo Microscope and immediately embedded in OCT (Tissue-Tek). For *Fos* expression (Figure 4B-F), mice were euthanized 2 hours after PE (10 mg/kg, i.p.) administration. Tissue blocks were cryosectioned at 10 μm, mounted onto Superfrost Plus slides (Thermo Fisher Scientific), and stored at −80 °C until use. Slides were fixed in freshly prepared 4% paraformaldehyde (PFA) in RNase-free PBS for 60 minutes at room temperature, followed by graded ethanol dehydration (50% to 100%). Sections were digested with protease IV for 30 minutes at room temperature, treated with HiplexUp reagent to quench endogenous fluorescence, and hybridized with the following probes: *Fos* (T4), *Ptger3* (T5), *Dbh* (T6), *Ddah1* (T7), *Vip* (T8), and *Npy* (T9). Imaging was performed in 4x SSC buffer using a Leica SP8 confocal microscope equipped with a 16x immersion objective (HC LUOTAR L 16×/0.8 IMM motCORR VISIR).

RNAscope Multiplex Assays (Figures 2B, S3D) were performed using the Multiplex Fluorescent Assay v2 kit (Advanced Cell Diagnostics) following the manufacturer’s protocol. Sample preparation and sectioning were conducted as described above. The following probe combinations were used: *tdTomato* (C1), *Ddah1* (C2) and *Npy* (C3). For images shown in Figure S3D, mCherry signal was detected via post-RNAscope immunostaining using goat anti-mCherry (1:600).

### Surgical delivery of AAVs and DT for ICN tracing, neuromodulation, and ablation

Surgical delivery of AAVs and DT offers a powerful approach for achieving spatially restricted genetic manipulation in the ICNS. For anatomical tracing and neuromodulation, AAV9-flex-tdTomato (Addgene #28306) and AAV9-hSyn-DIO-hM3D(Gq)-mCherry (Addgene #44361) were used. Both viruses contained 10^12^ - 10^13^ viral genomes per ml. For ablation studies, DT (Sigma D0564) was injected at a dose of 5 μg/kg. Fast Green FCF (0.05%, Sigma-Aldrich F7252) was occasionally included in the injection solution to facilitate visualization.

Mice were anesthetized with 1-2% isoflurane and maintained on a heated surgical platform. Analgesics were administered subcutaneously: meloxicam (5 mg/kg) and buprenorphine XR (Ethiqa, 3.25 mg/kg). Animals were intubated and mechanically ventilated using a mouse ventilator (Harvard Apparatus Model 687). A transverse sternotomy^45^ was performed to expose the heart: the sternum was carefully exposed and divided to gain access to the central atrial surface. AAVs, DT, or saline as control was injected (3.5 μl) directly into the atrial fat pad using a Nanoject III injector (Drummond). Mice were allowed to recover and monitored post-operatively. Tissues were harvested three weeks post-injection for histological analysis.

### Histology

Mice were anesthetized with 3-5% isoflurane and transcardially perfused with cold PBS (pH 7.4) containing 50 μg/ml of heparin (Sigma-Aldrich, H3149), followed by cold 4% PFA. Target tissues were post-fixed in 4% PFA at 4 °C (whole brain: overnight; half heart: 5 hours; whole heart: overnight; stellate ganglia: 4hours). For cryosectioning, tissues were cryoprotected in 30% sucrose in PBS for 1-3 days, embedded in OCT, and stored at −80 °C until use. For tissue clearing using modified CUBIC or iDISCO protocols^46,70,71^, samples were washed three times in PBS followed by three washes in PBS containing 0.01% sodium azide (NaN_3_; w/v; Sigma S8032) and stored at 4 °C until further processing.

#### Immunostaining on cryosections

Cryoprotected tissues were sectioned at 40 μm and mounted on Superfrost Plus slides (Thermo Fisher Scientific). For images shown in Figures 2A (Ddah1^tdT^ and Npy^tdT^) and S8C, sections were washed (3x PBS), permeabilized (1% Triton X-100, PBS, 10 minutes), blocked (4% Normal Donkey Serum, PBST (PBS, 0.05% Tween-20), 1 hour) and incubated with primary antibodies (Figure 2A: guinea pig anti-PGP9.5, 1:600 and rabbit anti-RFP, 1:500; Figure S8C: goat anti-mCherry, 1:600 and rabbit anti-cFos, 1:500) diluted in blocking buffer for 4 hours at room temperature or overnight at 4 °C. After washing (3x PBST), slides were incubated with corresponding Alexa Fluor 488- and 594-conjugated secondary antibodies (1:1,000, blocking solution, Jackson ImmunoResearch) for 2 hours at room temperature. Images showing reporter signals in the brain and stellate ganglia (Figure S3C and S3E-G) were from native fluorescence without immunostaining. Sections were mounted with ProLong Gold Antifade Mountant with DAPI (Thermo Fisher Scientific) and imaged using a Leica SP8 confocal microscope with 10x and 20x objectives.

#### Immunostaining on flattened wholemount atria

Wholemount heart atrial samples (Figures 2A (Vip^L10-GFP^ and Slc18a2^Sun1-GFP^), S3H and S5A) were cleared and stained as previously reported^69^ using a modified CUBIC protocol. Briefly, dissected tissues were first immersed into ½ water-diluted CUBIC Reagent-1 (w/w: 25% urea, 25% Quadrol, 15% Triton X-100) with gentle shaking at 37 °C for 3-6h, followed by incubation in undiluted Reagent-1 with shaking until tissues became optically transparent. Samples were then washed three times in PBS containing 0.01% NaN_3_, blocked in 2% Normal Donkey Serum (v/v; Jackson ImmunoResearch) in PBS with 0.1% Triton X-100 and 0.01% NaN_3_, and incubated with primary antibodies (Figure 2A, Vip^L10-GFP^: chicken anti-GFP, 1:500; Figure 2A, Slc18a2^Sun1-GFP^: chicken anti-GFP, 1:500, goat anti-PHOX2B, 1:600; Figure S3H: goat anti-mCherry, 1:600, guinea pig anti-PGP9.5, 1:600; Figure S5A-B: rabbit anti-RFP, 1:500) diluted in the same buffer for 7 days at room temperature with shaking. Following primary antibody incubation, tissues were thoroughly washed with PBS/0.1% Triton X-100/0.01% NaN_3_ and incubated with Alexa Fluor 488-, 594-, or 647-conjugated secondary antibodies in blocking buffer for 3 days at room temperature with shaking. After washing, samples were immersed in ½ PBS-diluted CUBIC Reagent-2 (w/w: 25% urea, 50% sucrose, 10% triethanolamine) overnight at room temperature, then transferred to undiluted Reagent-2 for 2 additional days. Finally, samples were stored in mineral oil (Sigma M5904) prior to imaging. For imaging, heart atrial tissues were mounted in a custom imaging chamber and gently flattened to ∼700 μm thickness. Cleared samples were imaged using a Leica SP8 confocal microscope with 10x or 20x objectives. Maximum-intensity projections were generated using LAX software or ImageJ.

#### Whole heart clearing, immunostaining, and lightsheet imaging

Whole hearts (Figures 3 and S5C-D) were processed using a modified iDISCO+ protocol for tissue clearing and immunolabeling. Dissected hearts were dehydrated through a graded methanol/ddH_2_O series (20%, 40%, 60%, 80% and 2× 100%) for 1hour each at room temperature, followed by incubation in 100% dichloromethane (DCM, Sigma-Aldrich, 34856) for 1 hour and then three 1-hour washes in 100% methanol. Samples were then bleached overnight in 5% hydrogen peroxide in methanol at 4 °C. Rehydration was performed through a reversed methanol/ddH_2_O gradient (80%, 60%, 40%, 20%) for 20 minutes each at room temperature. Samples were incubated twice in 5% DMSO, 0.3 M Glycine diluted in washing buffer (PBS with 0.1% Triton X-100, 0.05% Tween-20, 2 mg/l heparin, 0.2% NaN_3_) for 2hours each, followed by three 1-hour washes in washing buffer. Blocking was carried out in washing buffer containing 5% Normal Donkey Serum for 1 hour at room temperature. Samples were incubated with primary antibody (rabbit anti-RFP, 1:500) diluted in blocking buffer for 7-10 days at room temperature with gentle shaking. After extensive washing (2x 2 hours and overnight) in washing buffer, samples were incubated with Alexa Fluor 594-conjugated anti-rabbit secondary antibody (1:1,000 in washing buffer) for 5-7 days at room temperature. Following immunolabeling, samples were dehydrated through another methanol/ddH_2_O gradient (20%-80% then twice 100%) for 1 hour per step, incubated twice in 100% DCM for 1 hour each, and finally cleared in dibenzyl ether (DBE, Sigma 108014) until optically transparent. Samples were transferred to ethyl cinnamate (ECi, Sigma 112372) at least one night prior to imaging. Cleared whole hearts were imaged using an UltraMicroscope Blaze lightsheet microscope (Miltenyi Biotec) equipped with 1x, 4x and 12x immersive objectives in ECi-filled cuvette. Imaging was performed with a z-step size of 2 μm and a light sheet numerical aperture of 0.35 NA. Image acquisition was performed using Imspector Pro7.1 software.

#### Image processing and visualization

Whole heart images (Figures 3 and S5C-D) were processed using Imaris 10.1.1 (Bitplane). Three-dimensional rendering and visualization were performed using the Surface object to define anatomical regions (e.g., atria, ventricles, aorta, pulmonary veins, as shown in Figure 3B) and the Filament object to trace nerve fibers. Filament tracing and segmentation were conducted with the aid of the integrated machine learning tools. ICN cell bodies were manually annotated using the Spot object feature. The normalized fiber density (Figure 3D) was calculated by dividing the total filament area within each cardiac region by the region area and the total number of labeled ICNs per sample. The normalized intraganglionic fiber density (Figure 3F) was calculated by dividing the manually traced fiber area around ICN ganglia by the total number of labeled ICNs in each sample.

### *Ex vivo* retrograde Langendorff heart perfusion and functional measurements

*Ex vivo* mouse heart perfusion was performed using a custom-modified Langendorff apparatus to assess cardiac function under controlled conditions. Mice were anesthetized with 3–5% isoflurane and anticoagulated with heparin (10 mg/kg, i.p.) prior to heart extraction. Hearts were rapidly excised and immediately placed in ice-cold Krebs–Henseleit buffer (KHB) composed of (in mM): 112 NaCl, 4.7 KCl, 1.2 MgSO₄, 1.2 KH₂PO₄, 25 NaHCO₃, 11 glucose, 2 CaCl₂, and 0.5 EDTA, equilibrated with 95% O₂ and 5% CO₂ (pH 7.4). The aorta was cannulated and connected to the Langendorff system for retrograde perfusion at a constant pressure of 60 mmHg at 37 °C. Hearts were allowed to stabilize for 10–15 minutes prior to recording. For aortic root diameter measurements (Figure 6G-I), a pressure ramp ranging from 20 to 80 mmHg was applied in 10 mmHg increment.

Functional cardiac parameters were continuously recorded throughout perfusion. Surface ECG was recorded using paired electrodes placed near the right atrioventricular junction and in the perfusion bath, and signals were amplified using a P511 AC preamplifier (Grass Instruments). LVP and RVP were measured with solid-state pressure catheters (Transonic) inserted through the ventricular apex and amplified using the ADVantage PV System ADV500 (Transonic). Perfusion pressure was monitored via a fluid-filled pressure transducer (TSD104A, Biopac) connected to a DA100C transducer amplifier (Biopac). Coronary flow was measured using an in-line flow probe and TS420 perivascular flowmeter (Transonic). All analog signals were digitized at 1 kHz using the MP160 data acquisition system (Biopac) and analyzed offline. In parallel, heart contractions and aortic root diameter were recorded at 60 Hz using a high-speed camera (RWD) with a 500 nm green filter to block most blue light from optogenetic stimulation while preserving sufficient signal for visualization. Functional metrics, including heart rate, P and R waves, systolic and diastolic LVP/RVP, mean coronary flow, and peak pressure derivatives (ΔP/Δt) were quantified using custom Matlab scripts.

For optogenetic experiments (Figures 4 and 6), mCherry labeled ICN ganglia were visualized under a Leica M205FCA Fluorescence Stereo Microscope. A 200 μm core optic fiber (Thorlabs), coupled to a 470 nm diode laser (200 mW, Opto Engine), was positioned on top of the ganglia for focal illumination. Light stimulation was controlled via TTL pulses generated by the MP160 system (5 ms light pulses, 60–100 mW/mm^2^ intensity; for heart rate effects, 50 Hz, 10 – 20 s; for aortic root diameter measurements, 20 Hz, 20 s). Changes in heart rate and PR intervals were analyzed in 1 s bins for temporal dynamics (Figures 4H, S6A) and statistically comparisons were performed between the time point 5 s before stimulation onset and 10 s after stimulation onset (Figures 4I, S6C). Aortic root diameters were analyzed in 5 s intervals and reported both as percentage changes relative to 30 s baseline (Figure 6K) and as raw diameter values before and at the end of light stimulation (Figure 6L).

### In vivo physiological measurements in anesthetized and awake mice

*In vivo physiological measurements in awake mice.* Mice were pre-shaved around the neck region under 1-3% isoflurane anesthesia one week prior to recording. During each session, body temperature was first monitored using a rectal probe, and physiological parameters, including heart rate (HR), breathing rate (BrR), and oxygen saturation (SpO2%), were then measured using the MouseOX Plus Pulse Oximeter (STARR Life Sciences) with a small-size mouse collar sensor placed around the neck. Data were acquired using Mouse OX Plus software and analyzed with custom MATLAB scripts. For baseline activity recordings (Figures 5B, S7A, and S7D), each animal was recorded for 5 minutes per time point, and average values were plotted. For PE-induced baroreflex experiments (Figures 4A, 4K-L, S6D-G, S7B), heart rate was recorded for 5 minutes before and continuously for 20 minutes following i.p. injection of PE (10 mg/kg, Sigma P6126), with values averaged in 1-minute bins. Both normalized heart rate values relative to baseline (Figures 4L) and absolute heart rate changes (Figure S6D-G) were analyzed. The minimum normalized heart rate during the 20-minute post-PE period was used to quantify PE-induced bradycardia (Figure 4K).

#### ECG recording in awake, restrained mice

ECG was recorded from awake mice restrained in custom-built chambers constructed from acrylic paneling and stereolithography 3D printed components (Figure S8A). The floor of each chamber featured two copper electrode pads positioned beneath the front-right and rear-left paws to approximate a lead II configuration. A piece of filter paper was placed under the animal to prevent urine from short-circuiting the signals. Voltage differentials between the two electrodes were amplified using a P511 AC preamplifier (Grass Instruments) and digitized with a sampling rate of 10 kHz using the MP160 data acquisition system (Biopac). Signals were analyzed offline using custom MATLAB scripts. Animals were acclimated to the chamber for 20 minutes per day on three consecutive days before DT injection, and for 10 minutes immediately before recording sessions after DT administration. For each animal and time point, recordings lasted 20 minutes or until sudden cardiac death occurred. Periods when mice moved without paw contact on the electrodes were excluded from analysis. P and R wave peaks were detected from ECG recordings, and heart rate was calculated on a beat-to-beat basis, then averaged into 1-minute bins. In Ddah1^DTR-DT^ mice that experienced sudden death during restraint, the abrupt drop in heart rate was used as a temporal reference point to align physiological events across animals (Figure 7C).

#### In vivo physiological measurements in anesthetized mice

Mice were anesthetized with 1-2% isoflurane and maintained at 37 °C using a feedback-controlled heating platform. Surface ECGs were recorded using two electrodes placed under the right forelimb and left hindlimb to approximate a lead II configuration. Electrode cream was applied between the paws and electrodes to enhance conductivity. ECG signals were amplified using a P511 AC preamplifier (Grass Instruments). To measure carotid blood flow, a midline neck incision (∼1.5 cm) was made to expose the common carotid artery, which was gently isolated from surrounding tissues. A perivascular flow probe (0.5 PSB, Transonic) was positioned perpendicular to the artery and connected to a TS420 perivascular flowmeter (Transonic). ABP was recorded via a 1.2 F Scisense solid-state pressure catheter (Transonic) cannulated into the right common carotid artery, with signals amplified using the ADVantage PV system (Transonic). To measure LVP and RVP, mice were intubated via tracheotomy and mechanically ventilated using a mouse ventilator (Harvard Apparatus). A left thoracotomy was performed by cutting over the 4^th^ – 6^th^ costal cartilage to expose the heart. Two calibrated 1.2 F pressure catheters were inserted into the left and right ventricles via the apex and connected to the ADVantage PV system for signal amplification. All physiological signals (ECG, ABP, LVP, RVP, and carotid flow) were digitized at 1 kHz using the MP160 acquisition system (Biopac) and acquired with AcqKnowledge software. Offline analyses were performed using custom MATLAB scripts. From ECG recordings, P and R wave peaks were detected to calculate heart rate and PR intervals. From pressure recordings, systolic, diastolic, end-systolic, and end-diastolic peaks were extracted. Derived parameters including systolic and diastolic ABP, maximum LVP and RVP, heart rate and PR intervals, maximum and mean carotid blood flow, and integrated carotid blood flow per heart cycle were then computed. To assess aortic root diameter *in vivo*, measurements were taken immediately following open-chest heart injection and at the conclusion of each physiological recording session (Figure 6E-F). The aortic root was surgically exposed and carefully dissected free from the pulmonary artery. Diameter was measured using a digital caliper under a dissecting microscope, and the process was recorded using a digital camera (RWD). Absolute diameter values were plotted over time for both control and ablated groups.

### Atropine test

To assess whether blockade of muscarinic acetylcholine signaling could rescue physiological changes following ICN ablation, we administered atropine intraperitoneally. To confirm the effectiveness of atropine in antagonizing parasympathetic tone, wildtype mice were injected with atropine (0.5 mg/kg, i.p., Sigma Y0000878) or saline, followed by PE (10 mg/kg, i.p.). Heart rate responses were recorded using the collar sensor system as described above (Figure S7B). As shown, atropine effectively blocked PD-induced baroreflex bradycardia. To test its potential protective effect, Chat^DTR-DT^ and Chat^DTR-saline^ received atropine (0.5 mg/kg, i.p.) twice daily beginning one day after DT injection, for a total of five days. Awake physiological parameters were recorded daily using the collar sensor system (Figure S7D).

### Stress paradigms and survival analysis following ICN manipulation

Ddah1^DTR-DT^ and littermate control lox^DTR-DT^ mice were used to assess the effects of ICN ablation under various physiological stress conditions. All animals underwent DT injection as described above and were allowed to fully recover prior to stress testing. For the physical restrain test, mice were subjected to 20 minutes of daily restraint in custom-built chambers for up to 10 consecutive days. For the chronic heat stress test, mice remained in their home cages and were housed in a temperature-controlled incubator set at 37 °C for 7 consecutive days. For the sympathetic overactivation stress test^57^, mice received a single i.p. injection of epinephrine (10 mg/kg, Sigma E4250) and caffeine (120 mg/kg, Sigma C0750). In all paradigms, mice were closely monitored every 3-12 hours to assess survival, and Kaplan-Meier survival curves were generated accordingly (Figure 7B, 7F, and 7H).

#### Sympathetic overactivation and ECG recording

To evaluate cardiac responses to sympathetic overactivation, Ddah1^DTR-DT^ and lox^DTR-DT^ mice were anesthetized with 1-2% isoflurane, and body temperature was maintained at 37 °C using a feedback-loop controlled heating platform. Surface ECG was recorded as described above at 1 kHz. Following a 5-minute baseline recording, mice received a single i.p. injection of epinephrine (10 mg/kg) and caffeine (120 mg/kg) and were monitored continuously for up to 40 minutes to capture cardiac electrical activity.

#### Chemogenetic activation

Three weeks after viral delivery, Ddah1^AAV-hM3D(Gq)^ mice and their littermate controls (lox^AAV-hM3D(Gq)^) were injected with CNO (5 mg/kg, i.p., Abcam ab141704) once daily for three consecutive days. On the fourth day, all animals received a single i.p. injection of epinephrine (10 mg/kg, Sigma E4250) and caffeine (120 mg/kg, Sigma C0750). Daily CNO administration was then continued for up to three additional days (Figure 7J). Mice were monitored every 6-12 hours to assess survival, and Kaplan-Meier survival curves were generated for comparison between groups (Figure 7K). To confirm neuronal activation following chemogenetic stimulation, Chat^AAV-hM3D(Gq)^ mice were euthanized 30 minutes after CNO administration^72^ (5 mg/kg, i.p.) at three weeks post-viral injection. Hearts were collected for cFos immunostaining as described above (Figure S8C).

### Statistics and reproducibility

All statistical analyses were performed using GraphPad Prism 10. Data are presented as mean ± SEM unless otherwise specified. For pairwise comparisons, statistical significance was determined using two-tailed Student’s t test. For comparisons involving multiple groups and conditions, two-way ANOVA was performed followed by Šidák’s multiple comparison test. Kaplan-Meier survival curves were compared using the log-rank (Mantel-Cox) test. The specific statistical tests used are indicated in the corresponding figure legends.

## Notes

### Competing Interest Statement

The authors have declared no competing interest.

