## Supplementary figures and images for "Two Specialized Intrinsic Cardiac Neuron Types Safeguard Heart Homeostasis and Stress Resilience"

### Supplemental Figures

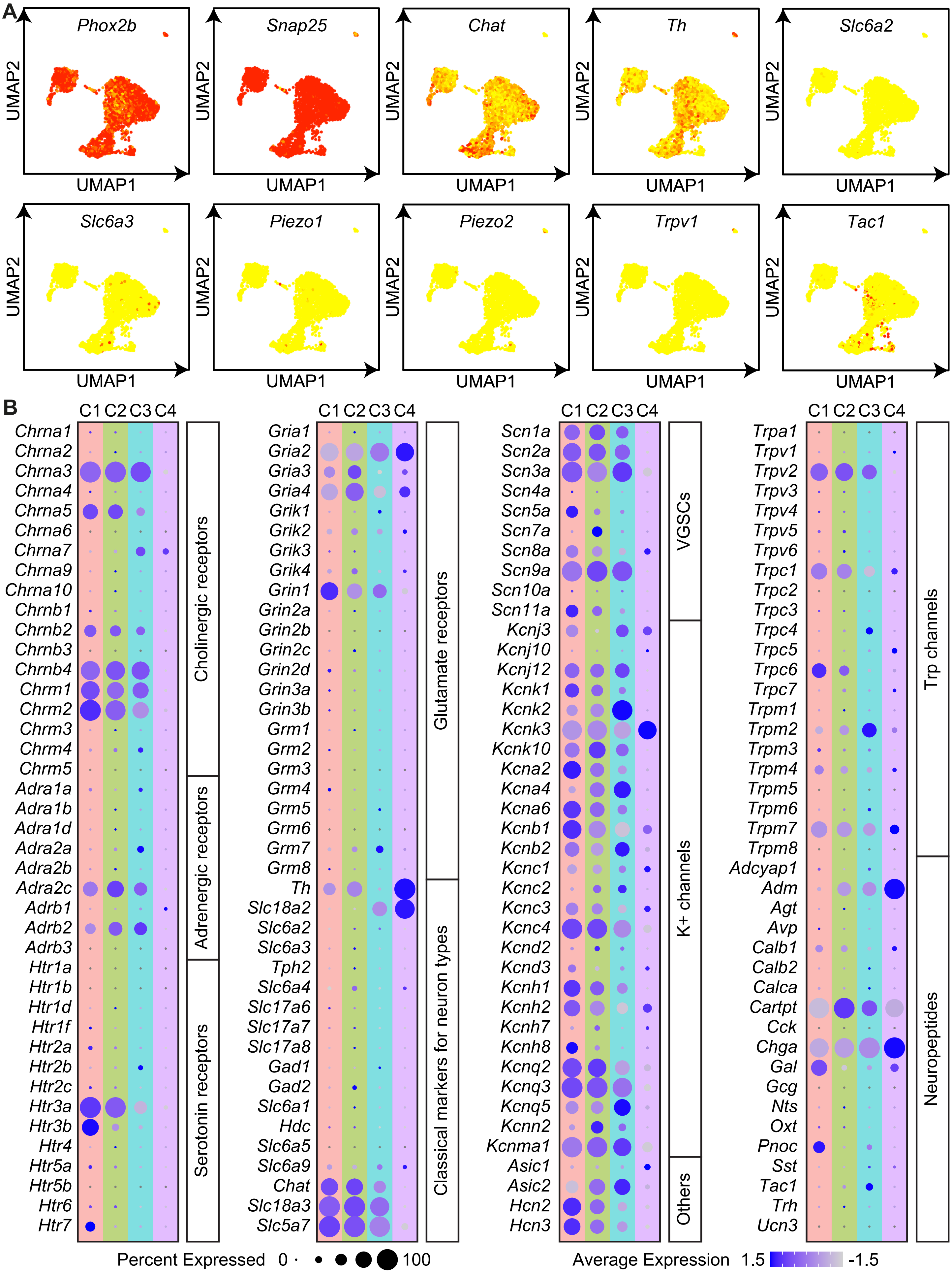

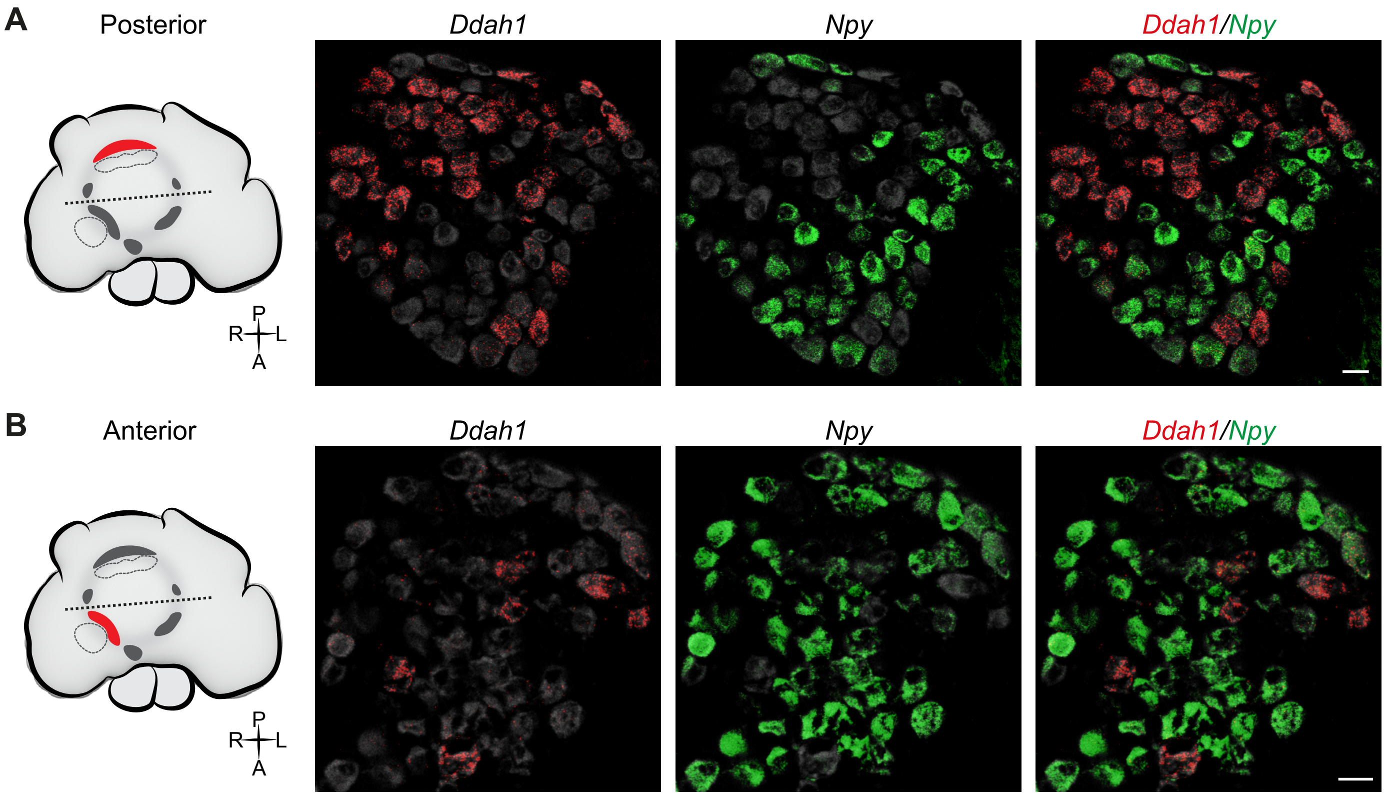

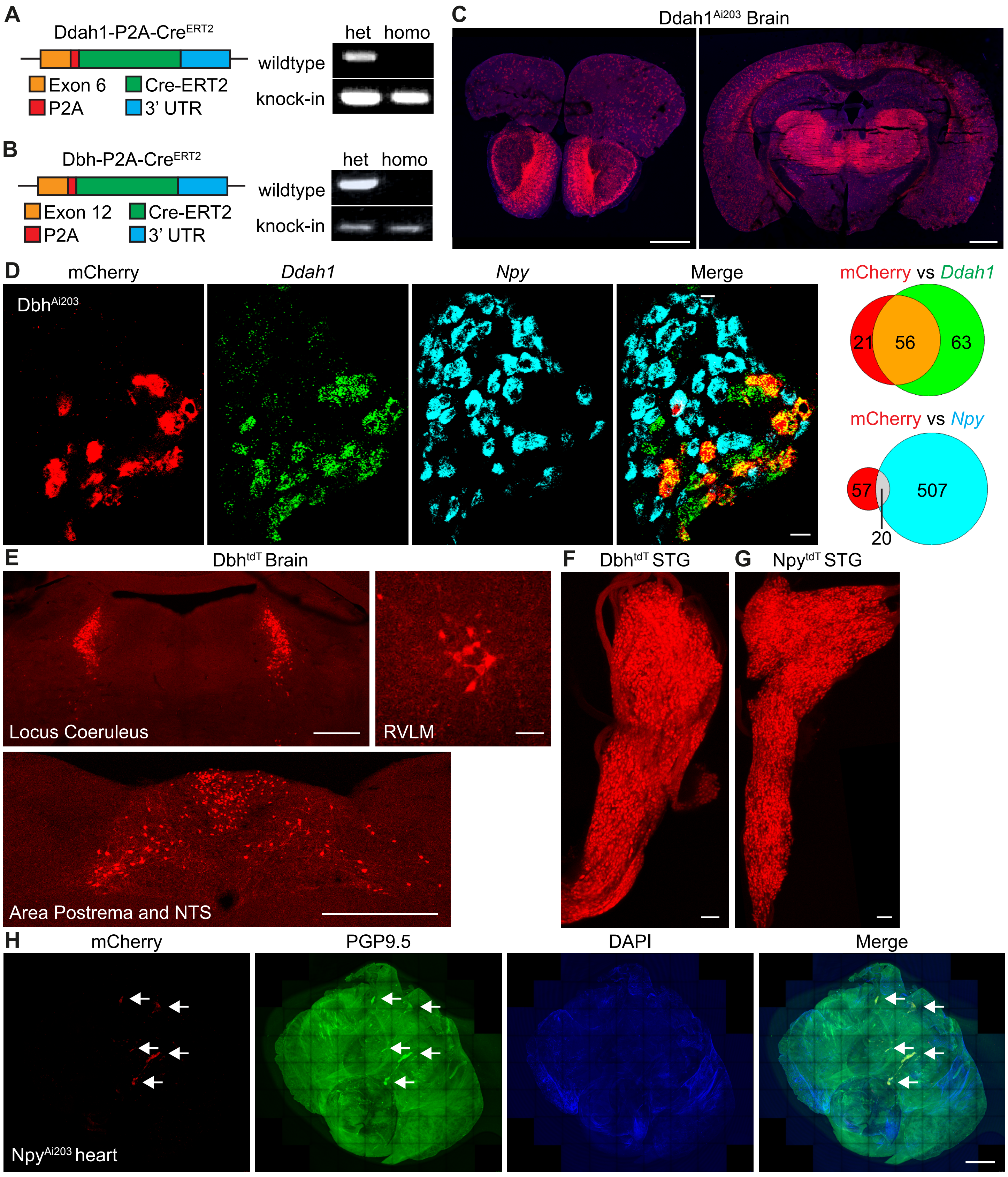

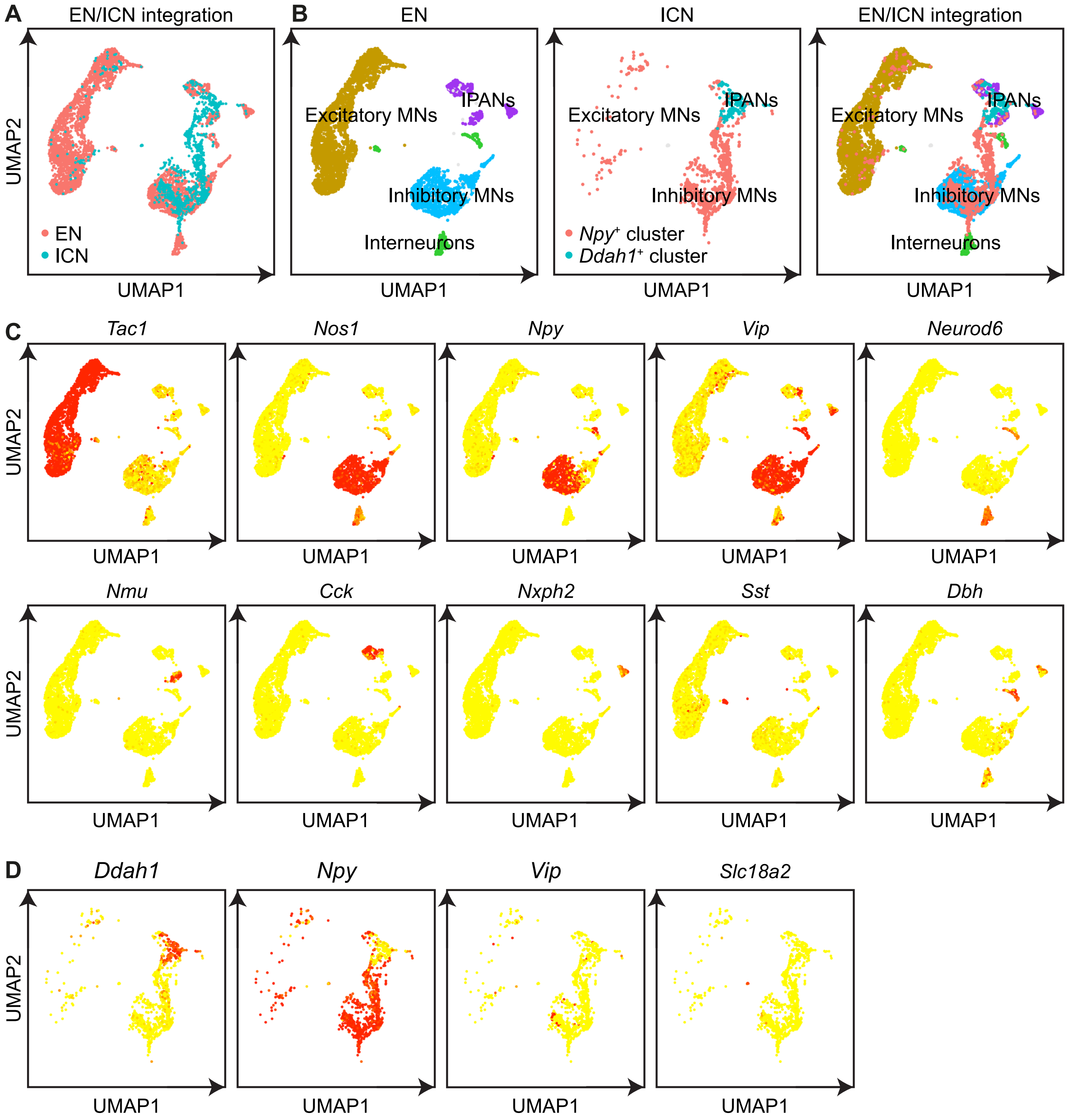

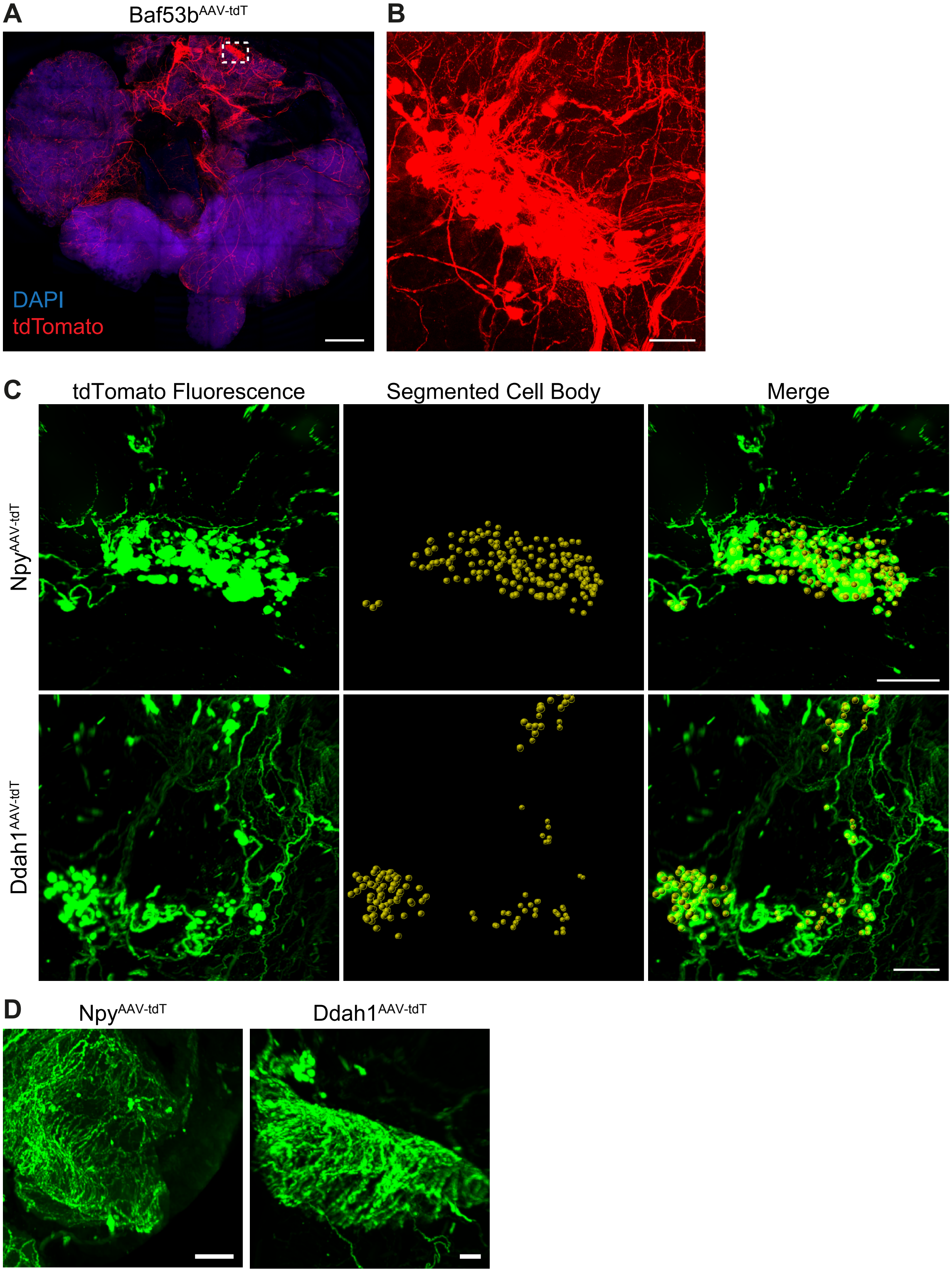

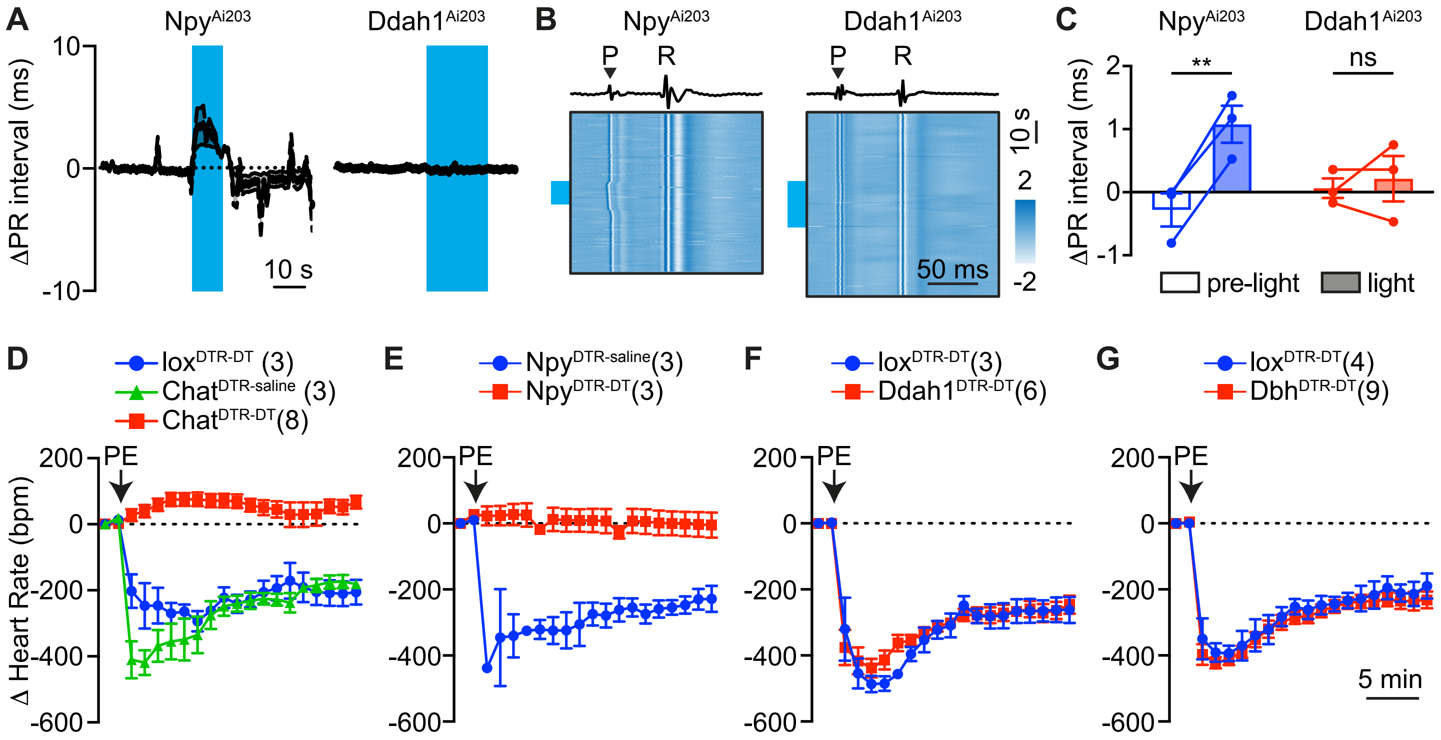

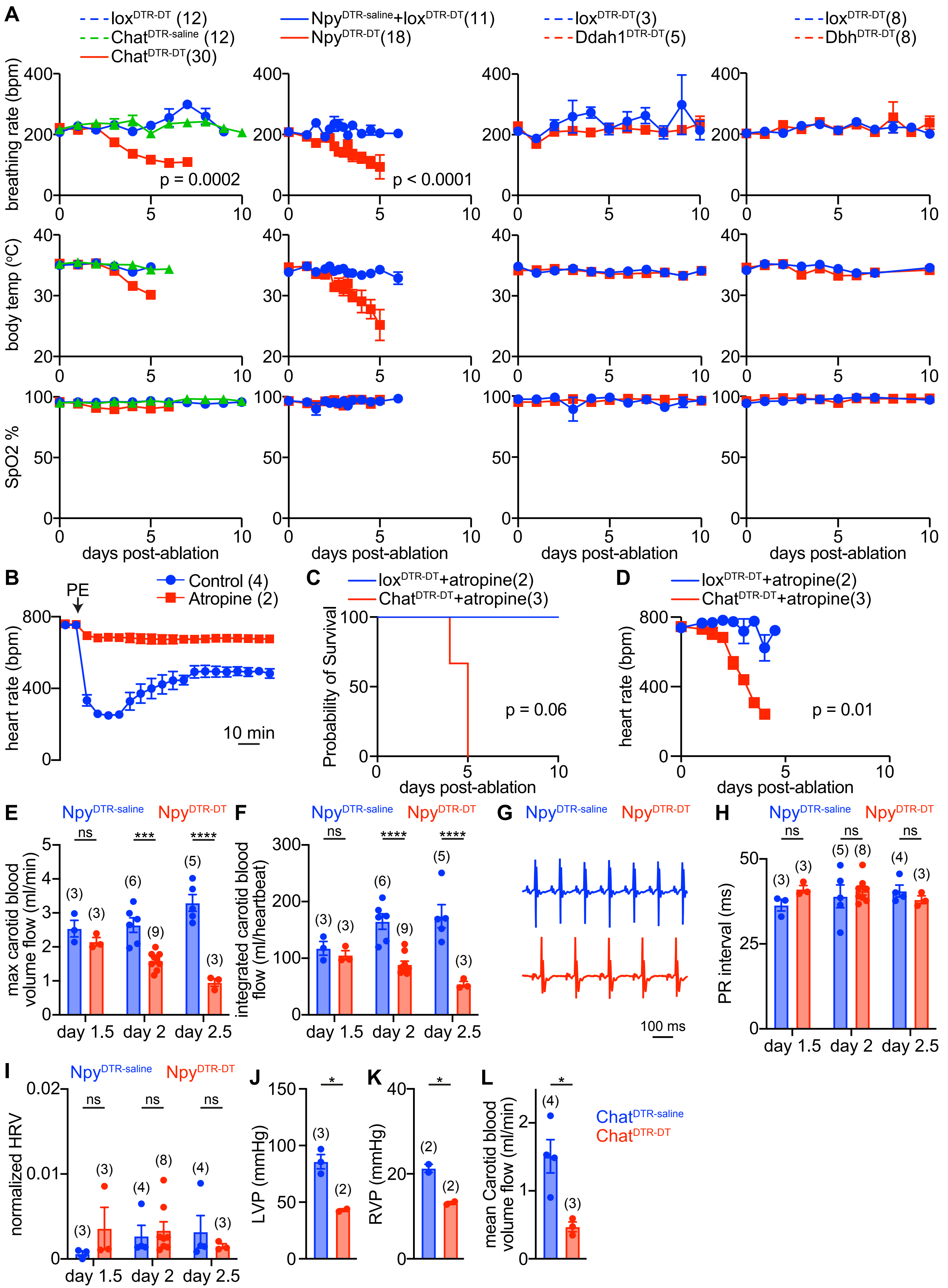

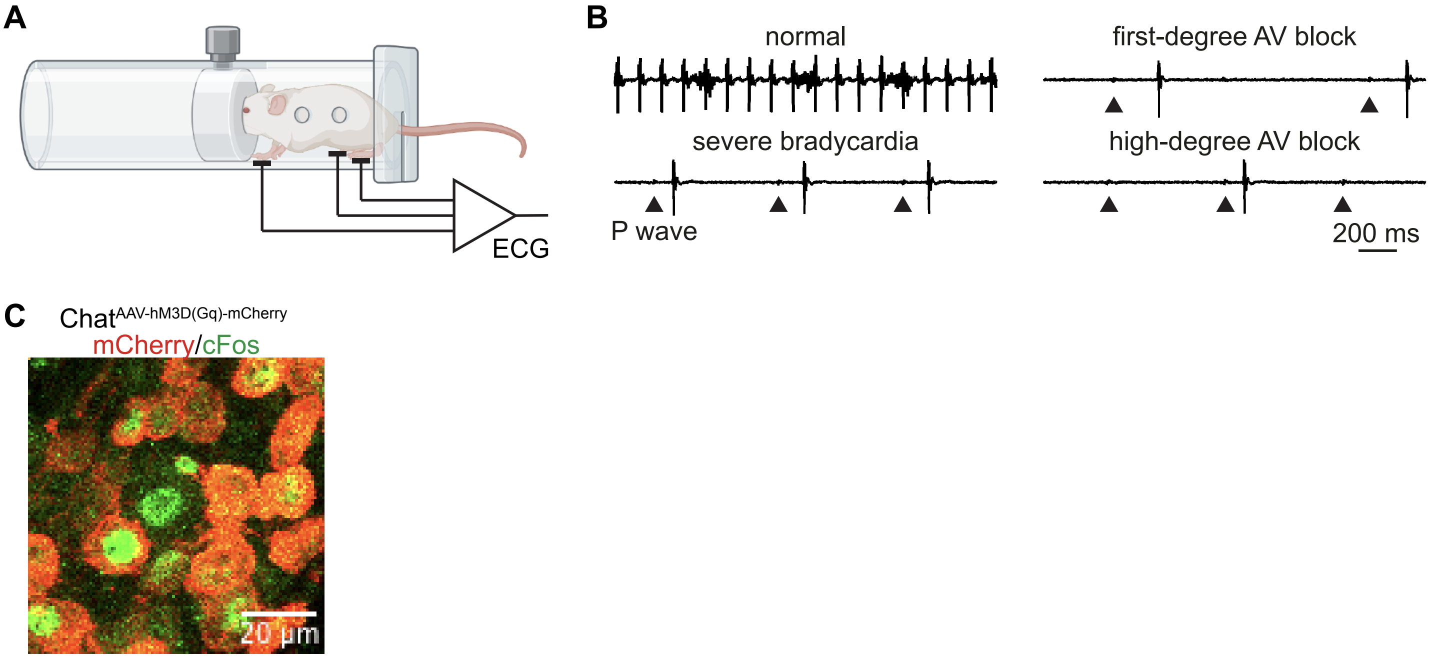
